# CD39^+^PD-1^+^CD8^+^ T cells mediate metastatic dormancy in breast cancer

**DOI:** 10.1101/2020.06.02.126037

**Authors:** Paulino Tallón de Lara, Héctor Castañón, Marijne Vermeer, Nicolás Núñez, Karina Silina, Bettina Sobottka, Joaquín Urdinez, Virginia Cecconi, Hideo Yagita, Farkhondeh Movahedian Attar, Stefanie Hiltbrunner, Isabelle Glarner, Holger Moch, Sònia Tugues, Burkhard Becher, Maries van den Broek

## Abstract

Some breast tumors metastasize aggressively whereas others remain in a state of metastatic dormancy for months or even years. The mechanism governing such metastatic dormancy remains largely unknown. Through high-parametric single-cell mapping in mice, we identified a discrete population of CD39^+^PD-1^+^CD8^+^ T cells present both in primary tumors and in dormant metastasis, which was hardly found in aggressively metastasizing tumors. Using blocking antibodies, we found that dormancy depended on TNF*α* and IFN*γ*. Of note, immunotherapy reduced the number of dormant cancer cells in the lungs. Adoptive transfer of purified CD39^+^PD-1^+^CD8^+^ T cells prevented metastatic outgrowth. In human breast cancer, the frequency of CD39^+^PD-1^+^CD8^+^ but not of total CD8^+^ T cells correlated with delayed metastatic relapse after resection (disease-free survival), thus underlining the biological relevance of CD39^+^PD-1^+^CD8^+^ T cells for controlling experimental and human breast cancer. Furthermore, density of CD39^+^PD-1^+^CD8^+^ T cells may serve as a novel biomarker and may serve as a potential immunotherapy target. Thus, we discovered that a primary breast tumor primes a systemic, CD39^+^PD-1^+^CD8^+^ T cell response that is essential for metastatic dormancy in the lungs.

## Introduction

Metastasis is the major cause of death in patients with breast cancer. The metastatic behavior of this disease is heterogeneous regarding affected organs as well as timing^1, 2^. Whereas some patients develop metastasis shortly after or even before diagnosis, others show metastatic lesions only years or decades after removal of the primary tumor^1, 3^. In fact, approximately 20% of disease-free patients will relapse within 7 years after resection, and patients that appear to be cured from breast cancer have a higher mortality than the rest of the population even 20 years after surgery^4, 5^. Such late relapses are thought to result from disseminated cancer cells (DCCs) that reached different organs but remained dormant for several years^6^. Indeed, the presence of DCCs in the bone marrow of breast cancer patients loosely correlates with the development of metastatic relapse^7, 8^.

The cancer cell-intrinsic and -extrinsic mechanisms governing metastatic dormancy are poorly understood. Some cancer cell-intrinsic factors have been associated with dormancy such as inhibition of the PI3K-AKT pathway^9^, activation of p38 or triggering of an endoplasmic reticulum stress response^10^. In addition, the microenvironment in the (pre)metastatic organ may induce dormancy^1^^1^ by different signals such as an endothelial cell-derived thrombospondin-1^12^ or TGF-β2 produced in the bone marrow^13^. Only few studies addressed the influence of cancer cell-extrinsic factors such as innate^14, 15^ and adaptive immunity^16, 17^ on dormancy. Although infiltration of the primary tumor by T cells was shown to correlate with a good prognosis^18^, it remains unclear whether this correlation is explained by cytotoxic T cells that eliminate cancer cells or by T cells that prevent the outgrowth of cancer cells and induce dormancy^19, 20^.

Here we sought to understand the mechanism that governs metastatic dormancy of DCCs and discovered that the primary tumor primes a systemic, CD8^+^ T cell response that prevents metastatic outgrowth of DCCs in the lungs. The protective T cells expressed markers that are generally associated with activation and exhaustion. Promoting such a response may provide a rationale for development of future immunotherapies that aim to prevent metastatic relapse.

## Results

### Disseminated 4T07 breast cancer cells are dormant

We used two preclinical models for breast cancer that were originally derived from the same spontaneous tumor in BALB/c mice, 4T1 and 4T07^21^ (Figure 1A). 4T1 orthotopic breast cancer produces macro-metastasis in different organs^22, 23^, whereas 4T07 is essentially non-metastatic. Some studies, however, showed the presence of disseminated 4T07 cells in distal organs^21, 24^. Orthotopically injected 4T1 and 4T07 cells showed comparable breast cancer growth and incidence in immunocompetent BALB/c mice, although 4T1 tumors grew faster at late stages (Supplemental Figures S1A and S1B). But, whereas 4T1 breast cancer readily induced aggressively growing and macroscopically visible lung metastasis, 4T07 did not (Supplemental Figures S1C and S1D).

**Figure 1.**
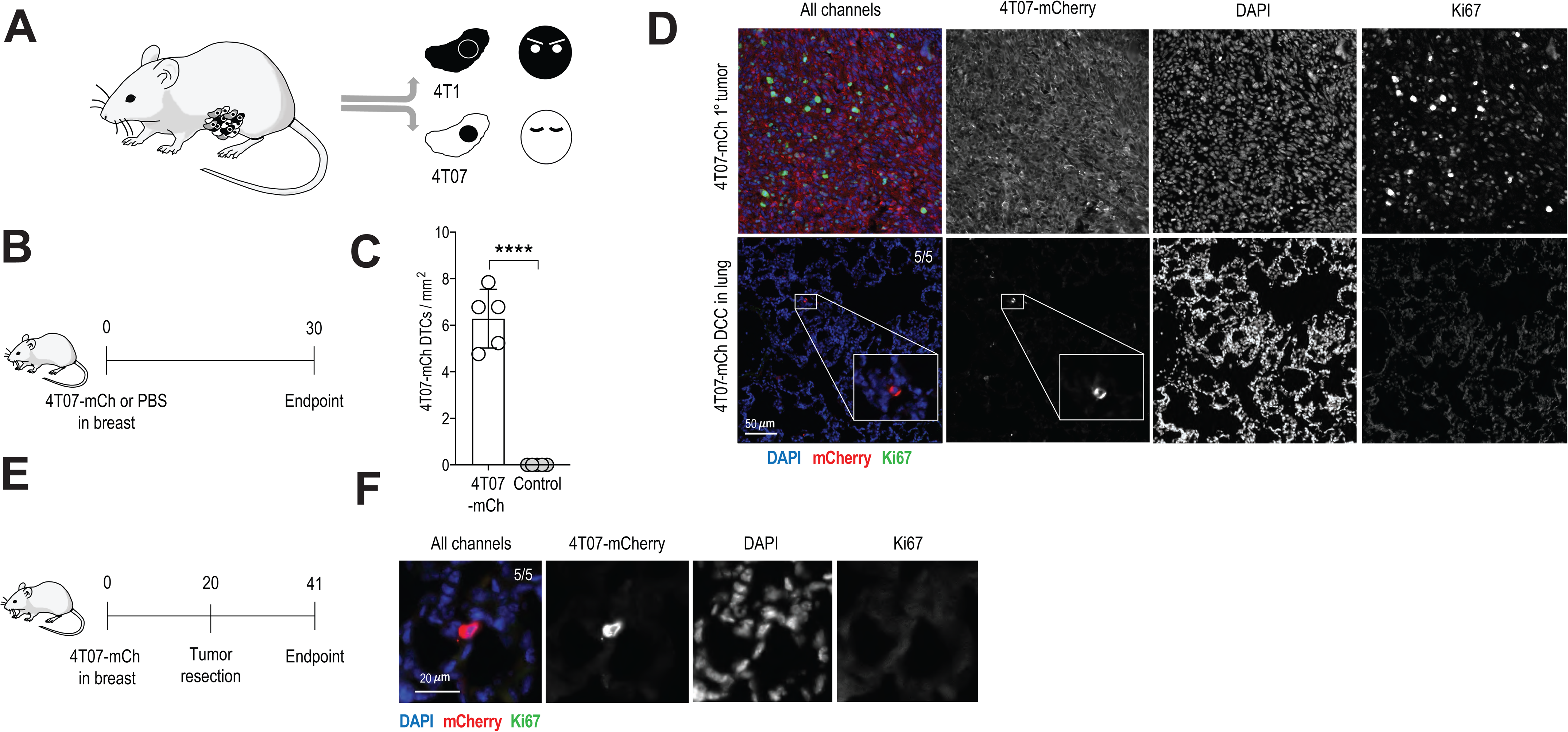
4T07 but not 4T1 breast cancer cells are dormant after dissemination to the lungs. **(A)** Diagram of the origin of 4T1 and 4T07 cell lines^21^. **(B)** Experimental design. 4T07-mCh cells (10^5^) or PBS were injected into the mammary fat pad of female BALB/c mice. Analysis was performed 30 d later. **(C)** Quantification of disseminated 4T07-mCh cells by quantitative pathology. **(D)** 4T07-mCh cells (10^5^) were injected into the mammary fat pad of female BALB/c mice. Analysis was performed 35 d later. Representative section of the primary tumor (upper panels) and lung (lower panels) at the endpoint. Disseminated 4T07 cells were detected as single, Ki67-negative cells in the lungs of 5 out of 5 mice. Disseminated 4T07 cells are shown in red, Ki67 in green, DAPI in blue. Scale bar indicates 50 µm. **(E)** Experimental design. 4T1 or 4T07 cells (10^5^) were injected into the mammary fat pad of female BALB/c mice. The tumor was resected 20 d after injection and analysis was performed 21 d after resection (endpoint). **(F)** Representative section of the lung at the endpoint. Disseminated 4T07 cells were detected as single, Ki67-negative cells in the lungs of 5 out of 5 mice. Disseminated 4T07 cells are shown in red, Ki67 in green, DAPI in blue. Scale bar indicates 20 µm. Each symbol represents an individual mouse. Five mice per group. \**p* < 0.05, \*\*\**p* < 0.001 (2-tailed Student’s t-test). The bar represents the mean ± SD. Results are representative of 3 independent experiments.

Despite absence of macro-metastasis, we found disseminated 4T07 cells in the lungs (Supplemental Figure S1E) of all mice, suggesting that 4T07 DCCs seed the lungs but fail to grow out. Presence of dormant DCCs in the lungs of all mice bearing a 4T07-mCh primary tumor was confirmed and quantified by immunofluorescence (Figures 1B, 1C and 1D, and Supplemental Figure S1F). Disseminated 4T07 cells were present as single cells and were Ki67-negative, as has been previously described for dormant DCCs^15, 17^. The fact that we found non-proliferating, disseminated 4T07 21 days after resection of the primary tumor (Figures 1E and 1F) further underscores the dormant state of 4T07 DCCs.

### Breast cancer induces CD8^+^ T cell-mediated immunity resulting in dormancy

We first investigated whether cancer cell-intrinsic features influenced the outgrowth of DCCs in the lungs by injecting 4T1 or 4T07 breast cancer cells intravenously (i.v.) into naïve mice (Figure 2A). Under these experimental conditions, both cell lines comparably formed macro-metastases (Figure 2B), suggesting that both cell lines can progressively grow in the lungs.

**Figure 2.**
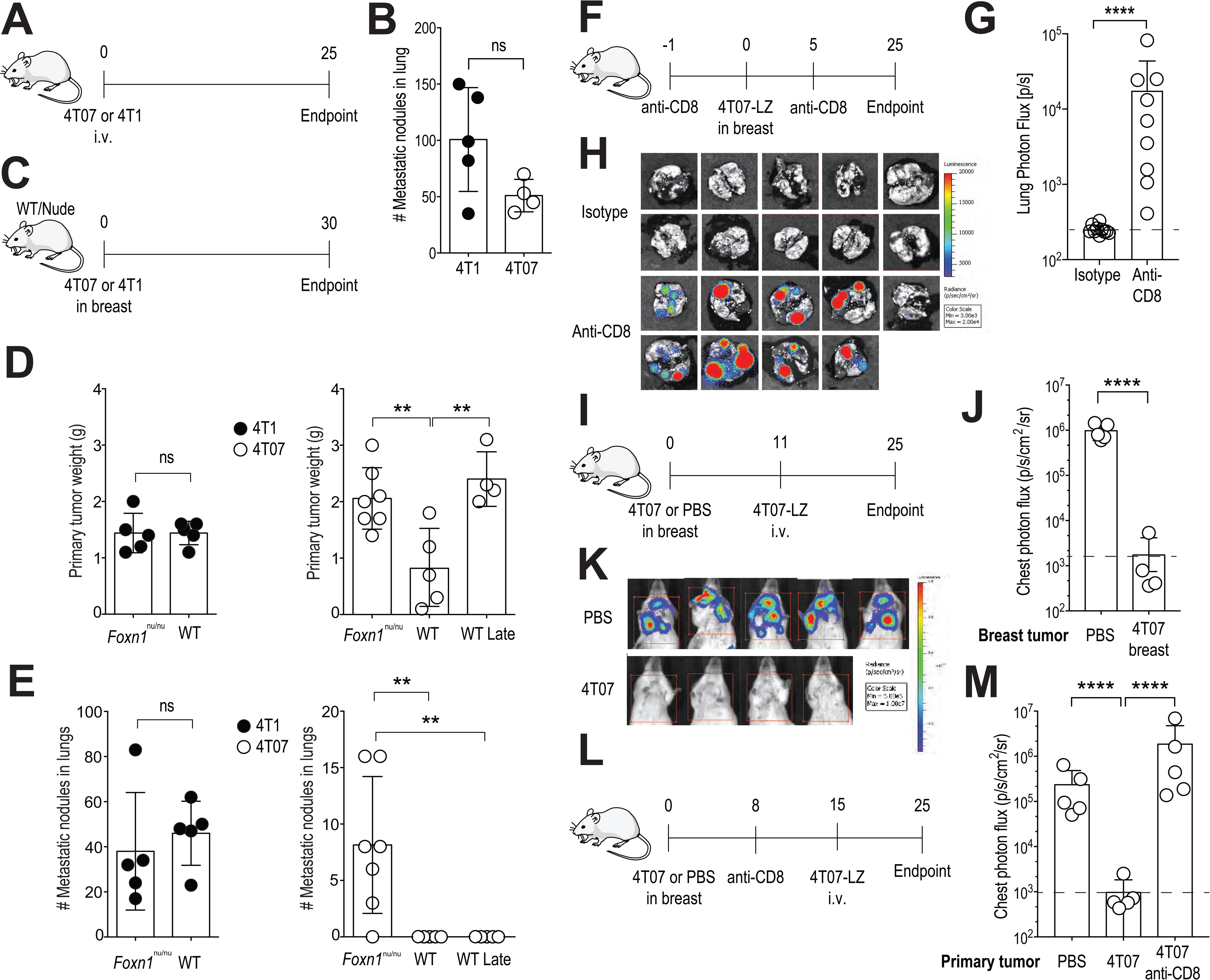
Induction of protective immunity by the primary tumor mediates metastatic dormancy. **(A)** Experimental design. 4T1 or 4T07 cells (3×10^5^) were injected i.v. into female BALB/c mice. Analysis was performed 25 d later. **(B)** Metastatic load in the lungs determined by counting nodules. **(C)** Experimental design. 4T1 or 4T07 cells (10^5^) were injected into the mammary fat pad of female BALB/c (*n*=5) or BALB/c *Foxn1^nu/nu^* (*n*=7) mice. Analysis was performed 30 d later. Lungs of “WT Late” mice (*n*=4) were analyzed when the 4T07 tumor size reached the size of the BALB/c *Foxn1^nu/nu^* group, e.g. 35 days after tumor cell injection. **(D)** Weight of primary tumors. Left panel with closed symbols, 4T1; right panel with open symbols, 4T07. **(E)** Metastatic load in the lungs determined by counting nodules. Left panel with closed symbols, 4T1; right panel with open symbols, 4T07. **(F)** Experimental design. Female BALB/c mice received 0.5 mg anti-CD8 or isotype control i.p. at days -1 and +5 relative to injection of 10^5^ 4T07-mCh cells in the mammary fat pad. Analysis was performed 25 d later. **(G)** Metastatic load in the lungs measured by bioluminescence. **(H)** Bioluminescence images. **(I)** Experimental design. 4T07 cells (10^5^) or PBS were injected into the mammary fat pad of female BALB/c mice on d 0. On d 11, 3×10^5^ 4T07-LZ cells were injected i.v. and analysis of lung metastatic load was performed on d 25. We chose d 11 because, on the one hand, this gives T cells enough time to be primed and expanded; on the other hand, it allows a long enough follow up after i.v. injection without the risk that the size of the primary tumor reached the humane endpoint of 225 mm^2^, which would result in forced termination of the experiment. **(J)** Quantification of lung metastatic load by bioluminescence. **(K)** Bioluminescence images. **(L)** 4T07 cells (10^5^) or PBS were injected into the mammary fat pad of female BALB/c mice on d 0. On day 8, mice were injected i.p. with 500 µg anti-CD8 or isotype control. On d 15, 3×10^5^ 4T07-LZ cells were i.v. injected and lung metastatic load was quantified on d 25. **(M)** Quantification of the lung metastatic load by bioluminescence on d 25. Each symbol represents an individual mouse. \*\**p* < 0.01, \*\*\*\**p* < 0.0001, ns = not significant (2-tailed Student’s t-test with Welch’s correction (B, D left panel, E left panel, G, J), ANOVA with Bonferroni correction (D right panel, E right panel, M). The bar represents the mean ± SD. Results are representative of at least 2 independent experiments.

The myeloid cell compartment was shown to undergo cancer-induced changes that promote metastasis by creating a pre-metastatic niche in the 4T1 model^25, 26^. Specifically, 4T1 primary tumors induce systemic neutrophilia^27^ and splenomegaly^26^, as well as accumulation of inflammatory monocytes^28^, eosinophils, neutrophils^27^ and alveolar macrophages^29^ in the pre-metastatic lung. We therefore compared 4T1- and 4T07-induced changes on the myeloid compartment in blood and lungs and observed no differences regarding splenomegaly (Supplemental Figure S2A), or neutrophilia (Supplemental Figure S2B). The lungs of mice with 4T1 or 4T07 breast cancer were similar concerning the presence of inflammatory monocytes (Supplemental Figure S2C), neutrophils (Supplemental Figure S2D), eosinophils (Supplemental Figure S2E) and alveolar macrophages (Supplemental Figure S2F) as determined by flow cytometry (Supplemental Figure S2G). We saw the previously reported accumulation of myeloid cells in the lungs during breast cancer progression^27^ in 4T1 and 4T07 alike (Supplemental Figure S2). Thus, the difference in metastatic behavior of 4T1 and 4T07 breast cancer cannot be explained by systemic changes in the myeloid compartment.

Orthotopic 4T07 tumors induced progressively growing lung metastases in T cell-deficient BALB/c *Foxn1^nu/nu^* mice, whereas T cell-deficiency hardly influenced metastatic behavior of 4T1 tumors (Figures 2C, 2D and 2E). To compare lung metastases in both strains, we analyzed wild type and *Foxn1^nu/nu^* mice at the same time point after injection and added an additional group of wild type mice in which breast tumors were allowed to progress until they matched the size in *Foxn1^nu/nu^* mice (WT late) (Figures 2D and 2E). Thus, metastatic dormancy of disseminated 4T07 breast cancer cells completely depends on T cells. To study whether CD8^+^ T cells are responsible for metastatic dormancy, we depleted CD8^+^ T cells from mice followed by orthotopic injection of 4T07 cells and subsequent analysis of lung metastatic load by IVIS (Figure 2F). While the growth of primary tumors was unaffected by CD8-depletion, disseminated 4T07-LZ cells grew out to macro-metastases in the absence of CD8^+^ T cells (Figures 2G and 2H), suggesting that primary 4T07 breast cancer induces CD8^+^ T cell-dependent immunity.

To test the hypothesis that CD8^+^ T cells are essential for metastatic dormancy, we orthotopically injected untagged 4T07 cells (or PBS as control) followed by an i.v. challenge with luciferase-tagged 4T07-LZ cells 11 days later (Figure 2I). If the primary tumor had induced protective immunity, we would expect metastatic dormancy of i.v. injected 4T07 cells in the lungs. Because we measured lung metastatic load by bioluminescence, we specifically quantified the i.v. injected, luciferase-tagged 4T07 cells. The presence of a primary 4T07 breast tumor prevented metastatic outgrowth of i.v. injected 4T07-LZ cells (Figures 2I, 2J and 2K) but did not influence the amount of seeding as measured 0.5 and 3 h after i.v. injection (Supplemental Figures S3A and S3B). At the endpoint, i.v. injected cells were present as disseminated, non-cycling single 4T07 cells in the lungs (Supplemental Figures S3C and S3D). Resection of the primary tumor before i.v. challenge did not interfere with dormancy. Specifically, i.v. injected 4T07-mCh cells readily induced macroscopically visible lung metastasis in control mice (PBS in breast and mock surgery; Supplemental Figures S3E and S3F), whereas only single, non-proliferating 4T07 cells were detected in the lungs of 4T07 tumor-bearing mice, independent of resection (Supplemental Figures S3E and S3G). We confirmed these results in a similar experimental set-up using 4T07-LZ cells and bioluminescence as read out (Supplemental Figures S3H and S3I). Depletion of CD8^+^ cells before i.v. challenge enabled metastatic outgrowth (Figures 2L and 2M), confirming the dependence of dormancy on CD8^+^ T cells. In contrast, 4T07 nor 4T1 breast cancer protected against experimental 4T1 lung metastasis, although 4T07-induced systemic immunity reduced the metastatic load resulting from i.v. injected 4T1-LZ cells (Supplemental Figures S3J and S3K).

Together, these data suggest that 4T07 breast cancer induces a systemic, protective immune response that mediates dormancy of DCCs in the lungs.

### CD39^+^PD-1^+^CD8^+^ T cells mediate metastatic dormancy

Since we found that 4T07 metastatic dormancy depends on CD8^+^ T cells, we characterized these cells in 4T1 and 4T07 primary tumors by high-dimensional flow cytometry. First, we gated on T cells (Supplemental Figure S4A), visualized all the markers in our 23-parameter panel using a two-dimensional *t*-stochastic neighbor embedding (tSNE) projection^30^ (Figure 3A) and clustered the cells using FlowSOM^31, 32^ (Figures 3A and 3B, upper panels and lower left panel). Based on the median marker intensities observed in the clusters (Figure 3B, lower right panel), we annotated the major T cell populations (CD8^+^ T cells, CD4^+^ T cells, and CD4^+^ CD25^+^ regulatory T cells) and determined their frequency and the position of each population in the tSNE projection. 4T07 primary tumors contained a higher proportion of CD8^+^ T cells (Figure 3B, lower left panel). We identified a discrete population of CD39^+^PD-1^+^CD8^+^ T cells present both in primary 4T07 tumors and in dormant metastasis, which was hardly found in aggressively metastasizing 4T1 tumors. The frequency of regulatory T cells was similar in 4T1 and 4T07 tumors, while conventional CD4^+^ cells were more abundant in 4T1 tumors compared to 4T07. Of note, within the CD8 compartment (Figure 3C), we observed that 4T07 breast cancer accumulated CD39^+^PD-1^+^CD8^+^ T cells (Figure 3D).

**Figure 3.**
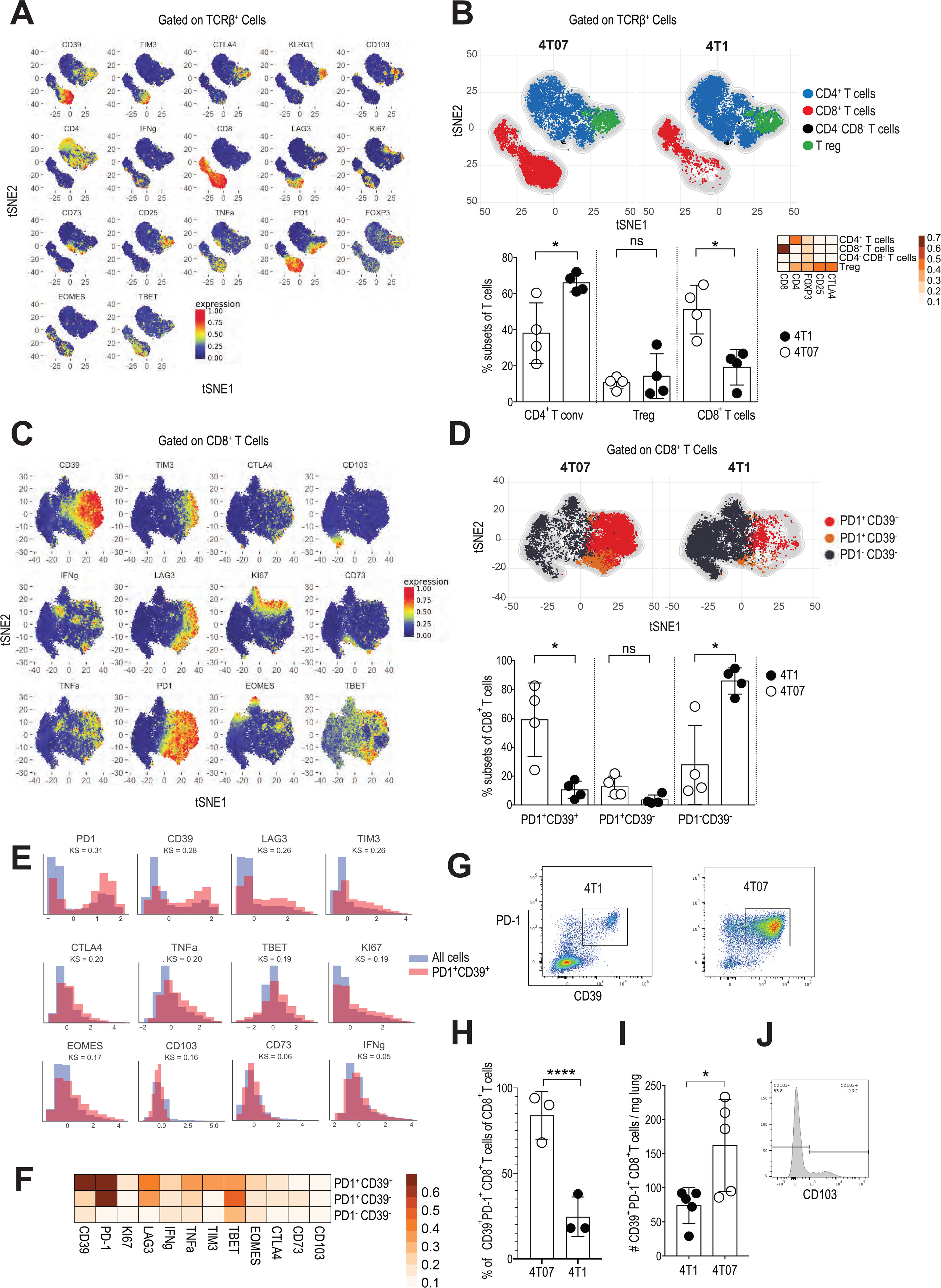
CD39^+^PD-1^+^CD8^+^ T cells emerge in dormant breast cancer. Female BALB/c mice were injected with 10^5^ 4T1 or 4T07 cells in the mammary fat pad and primary tumors were analyzed by flow cytometry 20 d later (*n*=4). **(A)** t-SNE visualization of markers after gating on single, live, CD45^+^ TCRβ^+^ CD44^+^ cells. **(B)** Upper panel: t-SNE plots of the main T cell subsets. Lower left panel: Frequency of the main T cell subsets in both groups. Each symbol represents an individual mouse. Lower right panel: Heat map with marker expression of the main T cell subsets. **(C)** t-SNE visualization of markers on CD8^+^ TCRβ^+^ cells. **(D)** Upper panel: t-SNE plots of the main CD8^+^ T cell subsets identified by FlowSOM algorithm. Lower panel: Frequency of the main CD8^+^ T subsets identified by FlowSOM algorithm in both groups. **(E)** Scaled histograms of arcsinh-transformed marker expression showing the relative marker distribution of the population identified by CellCnn (red) among all CD8^+^ T cells (blue). KS indicates the Kolmogorov–Smirnov two-sample test between the whole-cell population and the selected cell subsets. **(F)** Heat map with marker expression of the main CD8^+^ T cell subsets identified by FlowSOM algorithm. **(G)** Representative FACS plot of 4T1 and 4T07 primary tumors after gating on CD8^+^ TCRβ^+^. **(H)** Frequency of the population of total CD8^+^ identified by CellCnn in both models. **(I)** The number of CD39^+^PD-1^+^CD8^+^ T cells in the lungs. **(J)** Representative CD103-staining of live CD39^+^PD-1^+^CD8^+^ T cells from the lungs. Each symbol represents an individual mouse. \**p* < 0.05, \*\*\*\**p* < 0.0001 (2-tailed Student’s t-test with Welch’s correction). The bar represents the mean ± SD.

To substantiate that enrichment of CD39^+^PD-1^+^CD8^+^ T cells is the key immunological difference between 4T07 and 4T1 breast cancers, we analyzed our high-dimensional flow cytometry data using the unbiased representation-learning algorithm CellCnn^33, 34^. In agreement with FlowSOM analysis, CellCnn detected a population with high expression of CD39, PD-1, LAG3 and Tim-3 (Figure 3E and Supplemental Figure S4B). Indeed, the two markers that best defined the population and showed the biggest differential abundance in terms of the Kolmogorov–Smirnov two-sample test between the whole-cell population and the selected cell subsets (higher KS value) were PD-1 and CD39 (Figure 3E). This population was around 3 times more abundant in 4T07 compared to 4T1 tumors (Figure 3H, p<0.001).

In addition, CD39^+^PD-1^+^CD8^+^ T cells expressed more of LAG-3 and Tim-3 (Figure 3F), which characterizes this population as effector cells that experienced recent T cell receptor engagement^35, 36^ but may also indicate exhaustion^37, 38^. Therefore, we analyzed the functionality of CD39^+^PD-1^+^CD8^+^ T cells sorted from 4T07 tumors and confirmed the production of IFN*γ* and TNF*α* (Supplemental Figure S4C), suggesting that CD39^+^PD-1^+^CD8^+^ T cells in 4T07 tumors possess protective effector functions.

Importantly, CD39^+^PD-1^+^CD8^+^ T cells are found in the lungs of mice with primary 4T07 breast cancer (Figure 3I). A proportion of this population expresses CD103, hence resembling tissue-resident memory T cells^39, 40^ (Figure 3J). Our observation that CD39^+^PD-1^+^CD8^+^ T cells didn’t express CD103 in the primary tumor (Figure 3F) is in line with recent data showing that tissue-resident-like memory cells in human breast cancer are CD103-negative^41^. Thus, accumulation of CD39^+^PD-1^+^CD8^+^ T cells is the main immunological difference between dormant and metastatic tumors.

Because the CD8^+^ cells specifically emerging in 4T07 express PD-1, we wondered whether blocking PD-1 may improve their presumed protective effect. Indeed, immunotherapy with anti-PD-1 reduce the number of disseminated 4T07 cells in the lungs approximately twofold (Figures 4A and 4B), suggesting that the PD-1^+^ subset of CD8^+^ T cells comprises protective effector cells. To investigate whether the CD39^+^PD-1^+^CD8^+^ T cell subset indeed controls metastatic outgrowth and therefore mediates metastatic dormancy, we sorted this population from 4T07 breast cancer and adoptively transferred them to naïve BALB/c mice followed by an i.v. injection of 4T07-LZ cells (Figures 4C and 4D). Adoptive transfer of CD39^+^PD-1^+^CD8^+^ T cells prevented 4T07 lung metastasis, whereas injection of PBS or CD8^+^ T cells lacking CD39^+^PD-1^+^ cells (referred to as other CD8^+^ T cells) did not (Figures 4E and 4F). Thus, our results suggest that CD39^+^PD-1^+^CD8^+^ T cells control disseminated 4T07 breast cancer cells.

**Figure 4.**
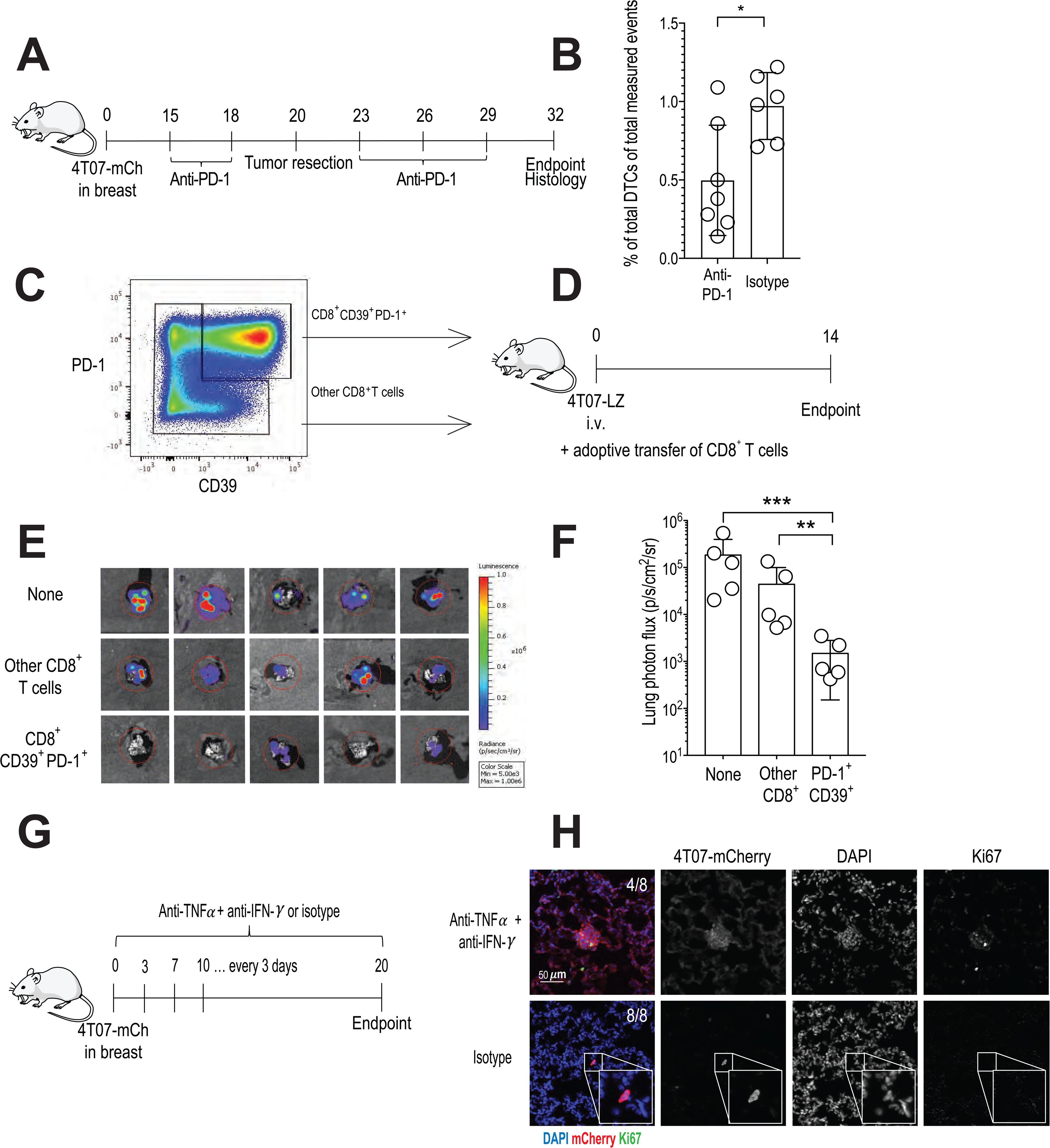
Tumor-associated CD39^+^PD-1^+^CD8^+^ T cells prevent metastatic outgrowth. **(A)** Experimental design. 4T07-mCh cells (10^5^) were injected into the mammary fat pad of female BALB/c mice. The breast tumor was resected on d 20 and histological analysis of lungs was performed on d 32. Mice received an i.p. injection with 250 µg anti-PD-1 or isotype control on d 15, 18, 23, 26 and 29. **(B)** Enumeration of disseminated 4T07 cells by quantitative pathology (right panel). Each symbol represents an individual mouse. \**p* < 0.05, (2-tailed Student’s t-test with Welch’s correction). The bar represents the mean ± SD. **(C)** Gating used for sorting of CD44^+^CD39^+^PD-1^+^CD8^+^ and the rest of the CD44^+^CD8^+^ population (termed other CD8^+^ T cells) from established 4T07 orthotopic breast tumors. **(D)** Experimental design. Two-hundred-thousand sorted CD44^+^CD39^+^PD-1^+^CD8^+^ or other CD44^+^CD8^+^ T cells were transferred after injection of 10^5^ 4T07-LZ cells into female BALB/c mice. All injections were given intravenously. Lung metastatic load was determined by bioluminescence 14 d later. **(E)** Bioluminescence of lungs at endpoint. **(F)** Quantification of lung metastatic load by bioluminescence. **(G)** Experimental design. 4T07-mCh cells (10^5^) were injected into the mammary fat pad of female BALB/c mice. Mice received an i.p. injection with 500 µg anti-IFN*γ* plus 500 µg anti-TNF*α* every 3^rd^ day. Control mice received isotype control antibody. On d 20, 4T07 cells were visualized in lungs by immunofluorescence. **(H)** Two representative examples showing clusters of proliferating 4T07 cells in the lungs of mice treated with anti-IFN*γ* plus anti-TNF*α*. Proliferating 4T07 cells were detected in the lungs from 4 out of 8 mice. Each symbol represents an individual mouse. Five mice per group. \*\**p* < 0.01, \*\*\**p* < 0.001 (ANOVA with Bonferroni’s correction). The bar represents the mean ± SD.

The presence of non-proliferating disseminated 4T07 cells in the lungs suggests that CD8^+^ T cells mediate dormancy by cell cycle arrest. Therefore, we tested whether CD39^+^PD-1^+^CD8^+^ T cell-derived IFN*γ* and TNF*α* can induce senescence^42^ by exposing 4T07 cells to those cytokines in vitro. We observed a significant increase in the number of senescent cells as measured by expression of senescence-associated *β*-galactosidase activity (Supplemental Figures S5A and S5B). To confirm the relevance of IFN*γ*/TNF*α*-induced senescence induction in vivo, we blocked both cytokines in mice with 4T07-mCh breast cancer and analyzed the status of disseminated 4T07 cells in the lungs by immunofluorescence (Figure 4G). Blocking IFN*γ* plus TNF*α* allowed the outgrowth of disseminated cells as illustrated by the presence of clusters of proliferating 4T07 cells (Figure 4H).

Thus, 4T07 primary tumors prime systemic CD8^+^ T cells that induce cell cycle arrest of DCCs by IFN*γ* and TNF*α* and thereby prevent overt metastatic disease.

### CD39^+^PD-1^+^CD8^+^ T cells have a transcriptome rich in effector and exhaustion molecules

To understand why this population controls DCCs, we sorted CD39^+^PD-1^+^CD8^+^ and other CD8^+^ T cells from 4T07 breast cancer and compared their transcriptome (Figures 5A and 5B and Supplemental Figure S6A). CD39^+^PD-1^+^CD8^+^ cells show overrepresentation of transcripts associated with effector function including *Prf1, Fasl, Gzmf, Gzme, Gzmd, Gzmc, Gzmb, Gzmk and Ifng* (Figure 5C). This was confirmed by gene set enrichment analysis (GSEA)^43^ showing that the transcriptome of the sorted CD39^+^PD-1^+^CD8^+^ cells shows strong similarities with human tissue-resident CD8^+^ T cells (Figure 5D), which were recently shown to correlate with improved survival in triple negative breast cancer^19^. This is in line with their unique capacity cells to prevent metastatic progression. Furthermore, the fact that dormancy is never observed in 4T1 breast cancer is explained by the low number of such protective T cells in 4T1-bearing hosts (Figure 3) and not by their transcriptional signature, since CD39^+^PD-1^+^CD8^+^ cells from 4T1 and 4T07 tumors did not show many differences (Supplemental Figures S6B, S6C and S6D). The transcriptome of CD39^+^PD-1^+^CD8^+^ cells was also enriched in markers of activation and exhaustion such as *Vsir* (VISTA), *Tnfrsf18* (GITR), *Tigit*, *Cd224* (2B4), *Tnfrsf9* (4-1BB), *Tnfrsf4* (OX-40), *Icos*, *Lag-3*, *Ctla4* and *Havcr2* (Tim-3). This is consistent with their overexpression of the transcription factor TOX (Figure 5C) which leads to expression of exhaustion markers and persistence of effector function during chronic antigen stimulation^44^.

**Figure 5.**
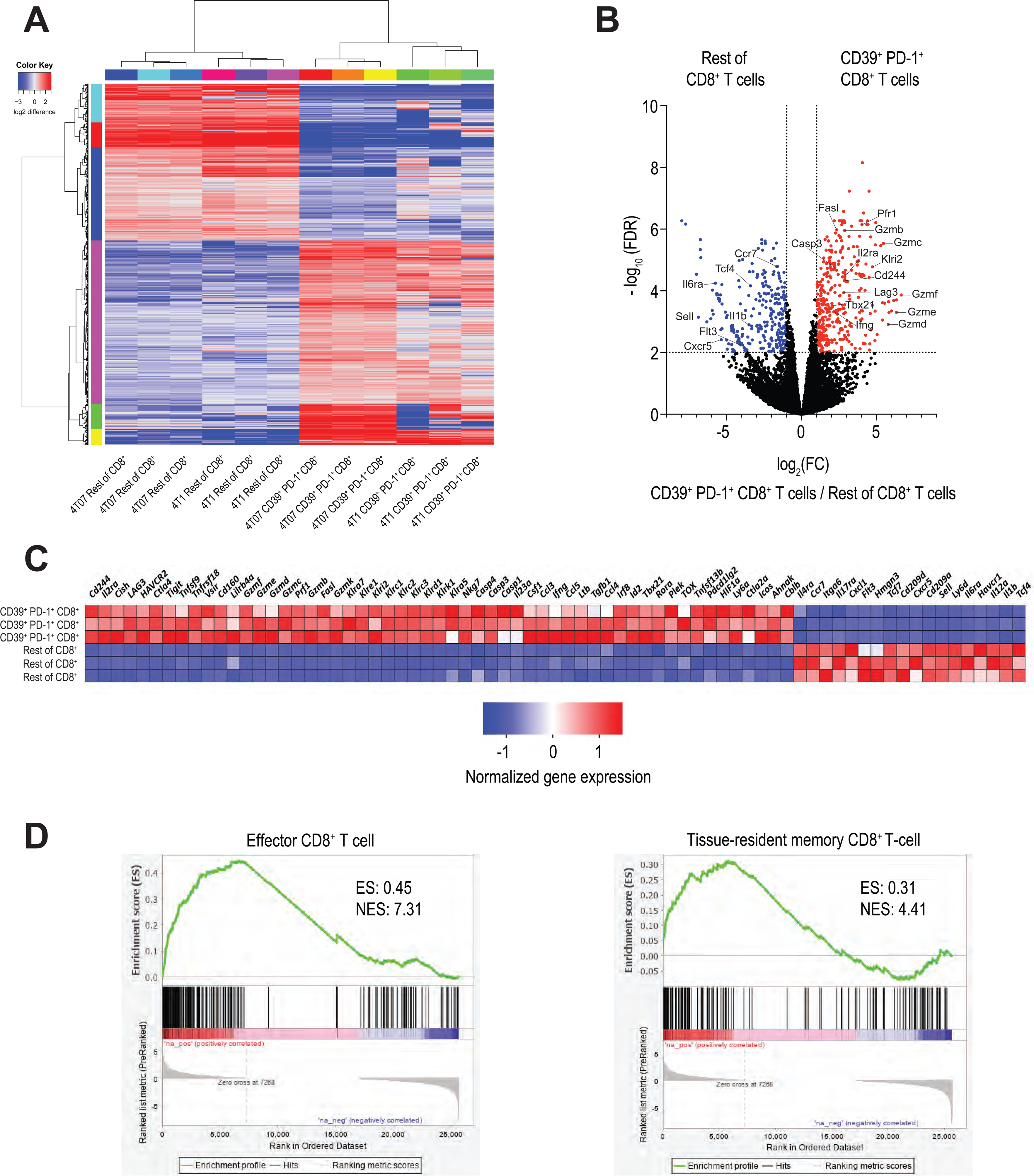
CD39^+^PD-1^+^CD8^+^ T cells have a unique transcriptional signature. **(A)** Hierarchical clustering of significantly differentially expressed genes comparing CD44^+^CD39^+^PD-1^+^CD8^+^ and rest of CD44^+^CD8^+^ T cells sorted from 4T1 and 4T07 breast tumors. **(B)** Volcano plot comparing transcripts in CD44^+^CD39^+^PD-1^+^CD8^+^ T cells with those in other CD44^+^CD8^+^ T cells sorted from 4T07 breast tumors. The red symbols represent transcripts that are significantly over-expressed in CD44^+^CD39^+^PD-1^+^CD8^+^ T cells, whereas the blue symbols represent significantly under-expressed transcripts. **(C)** Heat map showing relative expression of selected transcripts in CD44^+^CD39^+^PD-1^+^CD8^+^ (three top rows) and other CD44^+^CD8^+^ T cells (three bottom rows) identified by differential gene expression analysis. **(D)** Gene set enrichment analysis comparing the transcriptional profile to the data published by Goldrath et al.^78^ et al (left panel) and Savas et al.^19^ (right panel).

Taken together, CD39^+^PD-1^+^CD8^+^ cells have a unique transcriptomic signature defined by the co-expression of immune checkpoint molecules and effector proteins.

### CD39^+^PD-1^+^CD8^+^ but not total CD8^+^ T cells correlate with increased disease-free after resection survival in breast cancer patients

To investigate the clinical relevance of our preclinical findings, we analyzed primary breast cancer tissues from a cohort of 54 patients, who had a metastatic relapse after primary tumor resection (Table 1) for the presence of CD8, CD39, PD-1 and epithelial cells using 5-plex immunofluorescence (Figure 6A). Our cohort shows the typical survival curves for the individual subtypes, suggesting that it is representative (Supplemental Figure S7A). Patients with a high density of intra-tumoral CD39^+^PD-1^+^CD8^+^ T cells had a significantly longer disease-free survival after surgery than patients with a low density of such cells (Figure 6B). The density of extra-tumoral CD39^+^PD-1^+^CD8^+^ T cells did not correlate with disease-free survival (Supplemental Figure S7B), whereas to density of CD39^+^PD-1^+^CD8^+^ T cells independent of their location did (Supplemental Figure S7C). In addition, we observed similar data when focusing on luminal A and B patients (Supplemental Figures S7D and S7E). The density of intra-tumoral CD39^+^PD-1^+^CD8^+^ T cells was not an independent variable according to multivariate Cox regression analysis (Table 1). Importantly, the density of intra-tumoral CD8^+^ T cells showed no correlation with disease-free survival (Figure 6C), strongly supporting our preclinical observations, namely that that disseminated cancer cells are controlled by this specific subset of CD39^+^PD-1^+^CD8^+^ T cells.

**Figure 6.**
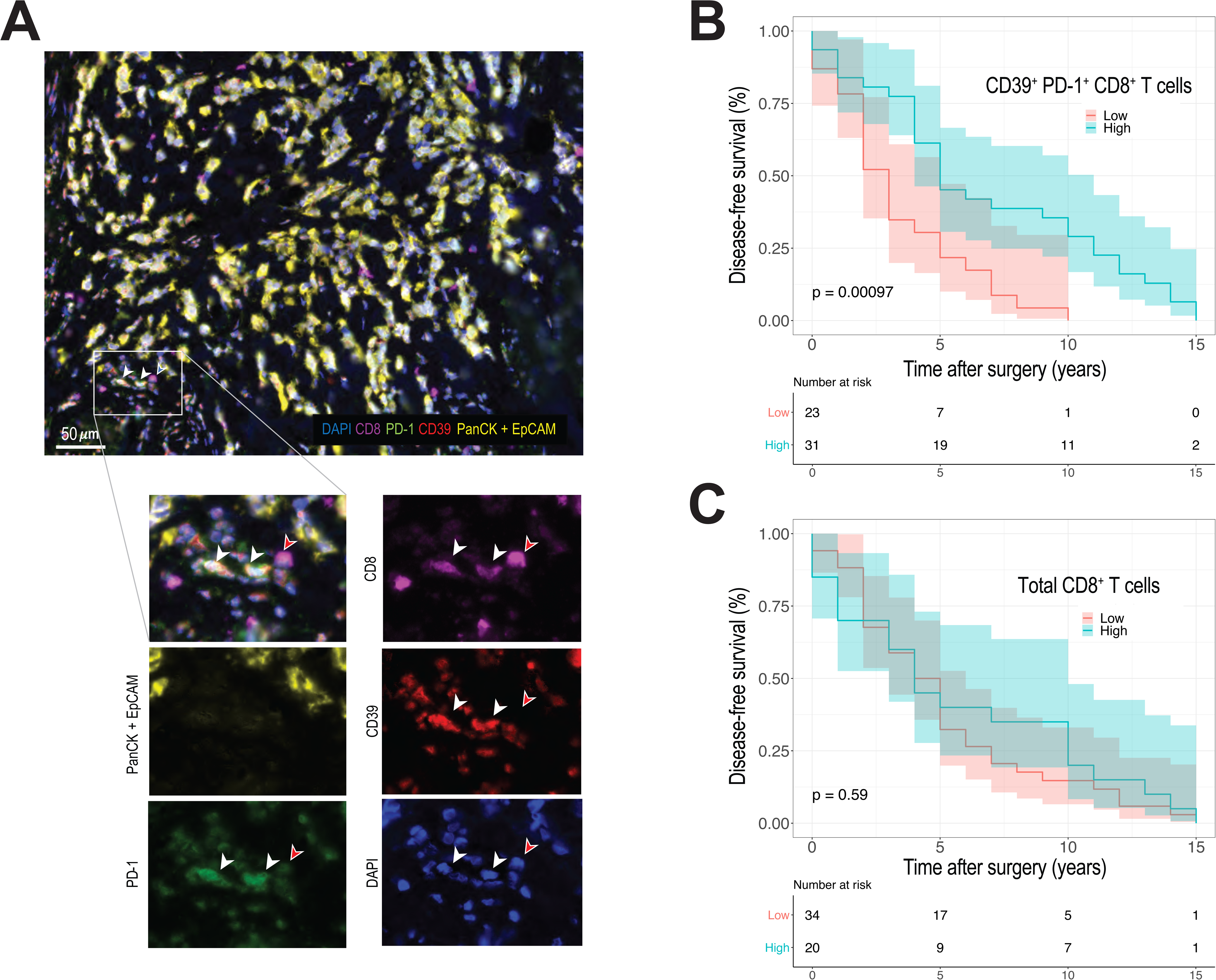
High density of intra-tumoral CD39^+^PD-1^+^CD8^+^ but not total CD8^+^ T cells correlates with disease-free survival in human breast cancer. **(A)** Representative images of 5-color multiplex immunofluorescence on human breast cancer. Staining shows epithelial cells (PanCK + EpCAM, yellow), CD8^+^ T cells (CD8, magenta), PD-1 (green), CD39 (red) and nuclear staining (DAPI, blue). Scale bar is 50 µm. **(B)** Disease-free survival of 54 patients with high or low number of intra-tumoral CD39^+^PD-1^+^CD8^+^ T cells**. (C)** Disease-free survival of 54 patients with high or low number of intra-tumoral CD8^+^ T cells. The threshold for separating patients with high and low CD39^+^PD-1^+^CD8^+^ T cell densities was defined using ROC curve analysis.

## Discussion

We discovered that breast cancer can prime a systemic, CD8^+^ T cell response that mediates metastatic dormancy in the lungs and consequently, protects against clinical metastatic outgrowth. We thus describe an unexpected trait of primary tumors, in addition to their well-known metastasis-promoting capacity^25, 26, 45, 46^. Using high-dimensional single-cell profiling of primary tumors, we identified the protective CD8^+^ T cell population as PD-1^+^ and CD39^+^ cells, suggesting recent cognate interaction and tumor-specificity^20, 47^. In line with our preclinical data, we found that a high density of intra-tumoral CD39^+^PD-1^+^CD8^+^ T cells significantly correlated with disease-free survival after resection in breast cancer patients. Even in patients with luminal A or B subtypes, where the tumor tissue is generally less infiltrated by CD8^+^ T cells^48^, the presence of CD39^+^PD-1^+^CD8^+^ T cells correlated with disease-free survival after primary tumor resection. As in our preclinical model, the heterogeneous population containing all CD8^+^ T cells did not correlate with disease-free survival, strongly supporting the notion that CD39^+^PD-1^+^CD8^+^ T cells comprise a population of cells that is uniquely equipped to control disseminated cancer cells.

The role of CD39 in cancer is rather complex. On the one hand, there is evidence that CD39 expression marks tumor-specific T-cells and presence of such cells correlates with a better prognosis or immune-mediated control^47^. On the other hand, CD39 is constitutively expressed by some cancer cells and suppressive cell types in the tumor microenvironment (Tregs and myeloid cells) and catalyzes pro-inflammatory extracellular ATP (eATP), resulting in impaired CD8^+^ T cell-dependent tumor control^49, 50^. The same papers described a reduction in tumor-associated macrophages and monocytes as well as a better T cell function (and tumor control) after blocking the enzymatic activity of CD39. We observed that CD39^+^PD-1^+^CD8^+^ T cells express effector molecules but also markers that are typically associated with exhaustion. A similar population was recently described in human breast cancer^19, 51–53^. Exhausted T cells are thought to have a reduced effector function and loss of proliferative capacity^54, 55^. Nonetheless, exhausted T cells in breast cancer may be more functional than in melanoma^56^.

In addition to local immunity, the induction of systemic immune responses by immunotherapy was shown to be essential for its efficacy^57^, suggesting that protective CD8^+^ T cells are recruited to the tumor. Recirculating CD8^+^ T cells can also enter tissues, where they develop into CD103^+^ tissue-resident memory cells and may protect tissues against disseminating tumor cells by the induction of cell death or cell cycle arrest^58^. Indeed, we detected CD39^+^PD-1^+^CD8^+^ T cells in the primary tumor and in the lungs and found that a proportion expressed CD103 as a marker of tissue residency. It was recently shown that the presence of intra-tumoral CD103-expressing CD8^+^ T cells correlate with survival in human breast cancer^41^.

As mentioned above, metastatic dormancy may result from two different states^59^: Disseminated single cells that are quiescent^60^, or micro-metastases that remain stable through an equilibrium between proliferation and death^61^. We found that disseminated 4T07 cells were present in the lungs as single non-cycling cells, suggesting dormancy through cell cycle arrest. Our observation that dormancy essentially depended on CD8^+^ T cells uncovers a novel mechanism underlying dormancy. We found that protective CD39^+^PD-1^+^CD8^+^ T cells produced TNF*α* and IFN*γ*, which were described to induce irreversible senescence in the context of Th1-mediated protection against cancer^42^; the same cytokines prevented proliferation of disseminated 4T07 breast cancer cells in the lungs.

It may be that environmental cues induce cancer stem cell-like properties in disseminated cancer cells^62^, which is in line with the low cycling frequency. Such cancer stem cells would represent a potent reservoir for progressive metastasis^63^ in response to environmental changes. Why dormant cancer cells sometimes awaken only after decades^3^ is still enigmatic, and it cannot be excluded that different pathways converge to initiate growth of dormant lesions. Assuming CD8^+^ T cells are required for maintenance of dormancy it is conceivable that attrition of tissue-resident memory CD8^+^ T cells may result in awakening of dormant cells. Tumor resection and consequent loss of antigen may result in diminished immunological memory; tissue-resident memory cells are described to be particularly unstable in the absence of antigen^64, 65^. Alternatively, dormant cells themselves may change, for example losing the expression of molecules preventing immune-mediated clearance^66^. Finally, environmental factors in the broadest sense may awaken dormant cancer cells. Such changes in the microenvironment include fibrosis^67^, tissue remodeling^68^, obesity^69^, inflammation^70^, disturbance of vascular homeostasis^71^, glucocorticoids^72^, or cigarette smoke^73^. How such factors induce cycling of dormant cells and whether immune cells can interfere at this stage is unknown. Here we discovered that CD39^+^PD-1^+^CD8^+^ T cells mediate dormancy of disseminated cancer cells but are by themselves unable to fully eradicate all of the cancer cells. This is clinically highly relevant, as dormant cancer cells are in all likelihood the source of future metastatic disease. Because the factors determining awakening are not known and may even not be controllable, it is important to better understand the nature of dormant cancer cells and which pathways must be mobilized for their complete elimination.

## Acknowledgments

We thank Anne Müller, Lubor Borsig and Christian Münz (all University of Zurich) for valuable input and support. We thank Fred Miller (Karmanos Cancer Institute) for providing cell lines. We thank the personnel from the Zurich Integrative Rodent Physiology (ZIRP, University of Zurich) and the Laboratory Animal Services Center (LASC, University of Zurich) for expert animal care. We thank Ruben Casanova (University of Zurich) for developing the R script converting immunofluorescence image data into file format compatible with flow cytometry data analysis software. This work was financially supported by the Swiss National Science Foundation (SNSF; CRSII5_177208 and 310030_175565 (MvdB), as well as 310030_170320, 310030_188450 and CRSII5_183478 (BB)), the Swiss Cancer League (Oncosuisse; KLS-4098-02-2017 (MvdB) and KFS-4431-02-2018 (BB)), the University of Zurich Forschungskredit (PTdL and HC), the University Research Priority Program (URPP) Translational Cancer Research (MvdB, BB), the Science Foundation for Oncology (SFO; (MvdB)), the Helmut-Horten Foundation (MvdB), the Hartmann-Müller Foundation Zurich (MvdB), the Monique-Dornonville-de-la-Cour Foundation Zurich (MvdB) and the European Union H2020 Project iPC #826121 (BB).

## Author contributions

PTdL, HC and MvdB conceived experiments; PTdL, HC, MV, NN, PC, EMC, VC, KS, FMA, JU, IG, MPL, SH performed and analyzed experiments; PTdL, HC, BB and MvdB wrote the manuscript; BB, ST, BS, HY and HM provided essential reagents; BB helped with data analysis; PTdL, HC and MvdB secured funding.

## Methods

### Mice

BALB/cJRj and BALB/cAnNRj*-Foxn1^nu/nu^* were purchased from Janvier labs (Roubaix, FR). NOD.Cg-*Prkdc^scid^Il2rg^tm1Wjl^*/SzJ mice were originally obtained from the Jackson Laboratory and provided by Christian Münz (University of Zurich, Switzerland). Mice were kept under specific pathogen-free conditions in individually ventilated cages at the Laboratory Animal Services Center at the University of Zurich. Mice had access to food and water *ad libitum* and were maintained on a 12-hour light/dark cycle with environmental enrichment. All experiments were performed with 8-14-weeks-old female mice in accordance with the Swiss federal and cantonal regulations on animal protection and were approved by The Cantonal Veterinary Office Zurich (156/2018).

### Cell lines

4T07 and 4T1 cells were a gift from Fred Miller (Karmanos Cancer Institute, Detroit, USA). Cells were cultured in Dulbecco’s Modified Eagle’s Medium (DMEM, Gibco) supplemented with 10% fetal bovine serum (FBS, ThermoFisher Scientific), 2 mM L-glutamine and 2% penicillin/streptomycin (ThermoFisher Scientific). Cells were cultured at 37°C in a humidified atmosphere with 5% CO_2_. 4T1 and 4T07 were lentivirally transduced to express firefly luciferase and ZsGreen (pHIV-Luc-ZsGreen, addgene plasmid #39196) or mCherry (pCDH-CMV-mCherry-T2A-Puro, addgene plasmid #72264). Transduced cells were sorted based on expression of GFP or mCherry, respectively. Luciferase-ZsGreen is referred to as LZ, mCherry as mCh.

Only cells of early passages were used for experiments. Cells were regularly tested negative for mycoplasma by PCR analysis. Cells were also tested negative for 18 additional mouse pathogens by PCR (IMPACT II Test, IDEXX Bioanalytics).

### In vivo tumor experiments and treatments

Hundred-thousand 4T1 or 4T07 cells in 50 µl PBS were injected into the fourth mammary fat pad. Alternatively, 3 x 10^5^ cells in 50 µl PBS were injected into the lateral tail vein.

For resection of primary tumors, mice were anesthetized with 2.5 % isoflurane and given 0.04 mg/kg fentanyl (Kantonsapotheke Zurich) i.p. as pre-emptive analgesia. Primary tumors were resected, and wounds were closed using Autoclip wound clips (BD Biosciences). For post-operative analgesia 0.1 mg/kg buprenorphine (Temgesic, Schering-Plough) was given i.p. immediately after surgery and in the drinking water at 10 µg/ml for 48 hours ad libitum.

For depletion of CD8^+^ T cells, 500 µg rat anti-mouse CD8 (clone YTS 169.4, hybridoma originally obtained from H. Waldmann, Oxford, United Kingdom) was injected intraperitoneally (i.p.) in PBS as described in each experiment. Control mice were injected with 500 µg rat anti-Trinitrophenol (clone 2A3, BioXCell). Antibodies were purified from hybridoma culture supernatant using protein G Sepharose 4 Fast Flow (Sigma). Injection of the respective antibody resulted in depletion of >90% of CD4^+^ or CD8^+^ T cells for at least 14 days, as determined by flow cytometry. Antibodies were administered i.p. in 200 µl PBS. Full depletion of the targeted population was confirmed by flow cytometry on blood 2 d after injection of the antibody in every experiment.

For blockade of PD-1, mice were injected with 250 µg anti-PD-1 (clone RMP1-14, made in-house by H. Yagita) as indicated. Control mice received 250 µg rat-anti-trinitrophenol (clone 2A3, BioXCell).

For blockade of INF*γ* and TNF*α*, mice were injected every 3^rd^ day starting on day 3 until the endpoint with 500 µg anti-INF*γ* (clone R4-6A2, BioXCell) plus 500 µg anti-TNF*α* (clone XT3.11, BioXCell) in 150 µl PBS. Control mice received 1000 µg rat-anti-trinitrophenol (clone 2A3, BioXCell).

Lung metastasis from luciferase- or mCherry-expressing tumors was quantified using an IVIS200 imaging system (PerkinElmer) as previously described^23^. Briefly, for luciferase-expressing tumors, mice were injected i.p. with 150 mg/kg D-luciferin (Promega) and photon flux was measured 20 minutes later in vivo as well as from dissected lungs. Lung metastasis from parental tumors was quantified by India Ink as previously described^23^. Briefly, mice were euthanized and India Ink (Pelikan, 15% in PBS) was injected intratracheally, lungs were harvested, washed in PBS and fixed in Fekete’s solution (62% ethanol, 3.3% formaldehyde, 0.25 M acetic acid). Metastatic foci were counted blinded using a dissection microscope.

Disseminated cancer cells were detected in the lungs using a colony forming assay as previously described^21^. Briefly, lungs were resected from euthanized mice, cut into small pieces and digested for 45 minutes at 37°C in DMEM with 1 mg/ml collagenase IV and 2.6 µg/ml DNase I (both Sigma) on a rotating device. Subsequently, samples were washed with PBS by centrifugation at 350 *g*. For detection of circulating cancer cells, blood was collected by heart puncture in a 25-gauge syringe containing 100 μl heparin (5’000 IU/ml, Braun) and cells were washed with PBS by centrifugation at 350 *g*. Blood and lung samples were suspended in complete DMEM containing 6 µM of 6-thioguanine and cultured in a T175 flask. Medium was exchanged after 5-7 days and presence of colonies of tumor cells was evaluated after 14 days by light microscopy and crystal violet staining.

### Immunofluorescence of mouse samples

Organs were fixed with 4% paraformaldehyde (Roti-Histofix 4%, Roth) for 10 minutes at RT and cryoembedded in Optimal Cutting Temperature (O.C.T.) Compound (O.C.T.^TM^ Compound, Tissue-Tek) using dry ice/100% ethanol slurry. Ten-µm thick sections cut using a cryotome (Leica), mounted on SuperFrost glass slides, dried for 1h at 37°C and preserved at -80°C until immunostained. To prevent unspecific binding of antibodies, slides were incubated with 4% donkey serum in PBS (ANAWA BioWorld) for 10 minutes at RT. Subsequently, slides were incubated overnight at 4°C with primary antibody diluted in PBS 1% donkey serum. Following primary antibodies were used: Anti-mCherry (goat polyclonal antibody, AB8181-200 SICGEN, 1:400), Ki67 (rabbit monoclonal antibody, ab16667 abcam, 1:100). After incubation, slides were washed 3 times with PBS Tween 20 0.05% and incubated for 1 h at room temperature with secondary antibodies diluted in PBS 1% donkey serum. Following secondary antibodies were used: AF594-conjugated anti-goat IgG, AF488-conjugated anti-rabbit IgG (both Jackson ImmunoResearch, 1:400). Finally, slides were washed, incubated with 0.5 µg/ml 4′,6 diamidine-2-phenylindole (DAPI; Invitrogen) for 5 min, washed again and mounted with ProlongDiamond medium (Invitrogen). The slides were scanned using the automated multispectral microscopy system Vectra 3.0 (PerkinElmer). An unstained slide was used to generate the spectral profile of autofluorescence in studied tissues. The Inform software (PerkinElmer) was used for spectral unmixing of individual fluorophores and autofluorescence.

### Flow cytometry

Animals were euthanized by isoflurane overdose. The right heart ventricle was perfused with 10 ml PBS to eliminate the blood from the lung vessels. Primary tumors and lungs were collected in DMEM, cut into small pieces and digested for 45 minutes at 37°C in DMEM containing 1 mg/ml collagenase IV and 2.6 µg/ml DNase I (both Sigma) on a rotating device. Samples were washed with PBS by centrifugation for 5 minutes at 350 *g*, the pellet was suspended in PBS and filtered through a 70-μm filter (BD Biosciences) to obtain a single cell suspension. For lymphocyte analysis, cells were further purified by centrifugation over a Percoll gradient (GE Healthcare, 17-0891-01, Sigma Aldrich).

Single cells were stained according to standard protocols. Briefly, cells were surface-stained in 50 µl antibody-mix in PBS. For intracellular cytokine staining, cells were stimulated with 100 ng/ml phorbol 12-myristate 13-acetate (PMA) plus 1 µg/ml ionomycin for 4 h at 37°C in the presence of GolgiPlug/GolgiStop (BD Pharmigen). Cells were stained for surface molecules as described above, washed with PBS, and fixed for 30 min on ice using IC Fixation Buffer from Foxp3/Transcription Factor Staining Buffer Set (eBioscience). Subsequently, cells were stained for intracellular cytokines in permeabilization buffer from the Foxp3/Transcription Factor Staining Buffer Set overnight at 4°C. After washing with permeabilization buffer, samples were suspended in FACS buffer (PBS, 20 mM EDTA pH 8.0, 2% FCS) and acquired using a CyAn ADP9 flow cytometer (Beckman Coulter), FACS LSRII Fortessa or FACSymphony (both BD Biosciences). For quantitative analysis, CountBright absolute counting beads were used (ThermoFisher Scientific). In all staining, dead cells were excluded using Live/Dead fixable staining reagents (Invitrogen), and doublets were excluded by FSC-A versus FSC-H and SSC-A versus SSC-H gating. Following directly labelled anti-mouse primary antibodies were used: Anti-CD8a in BUV 805, clone 53-6.7, rat IgG2a, BD. Anti-CD11b in BUV 661, clone M1/70, rat IgG2b, BD. Anti-CD45.2 in BUV 653, clone 30-F11, rat IgG2b, BD. Anti-VISTA in AF488, clone MH5A, armenian hamster IgG1, BioLegend. Anti-CD39 in PerCP-eFluor710, clone 24DMS1, rat IgG2b, ThermoFisher Scientific. Anti-LAG3 in BV 421, clone C9B7W, rat IgG1, BioLegend. Anti-CD44 in BV 570, clone IM7, rat IgG2b, BioLegend. Anti-CD73 in BV 605, clone TY/11.8, rat IgG1, BioLegend. Anti-CD25 in BV 650, clone PC61, rat IgG1, BioLegend. Anti-PD-1 in BV 785, clone 29F.1A12, rat IgG2a, BioLegend. Anti-TCR*β* in PE-Cy5, clone H57-597, armenian hamster IgG1, BioLegend. Anti-KLRG1 in APC-Cy7, clone 2F1/KLRG1, armenian hamster IgG1, BioLegend. Anti-TIM3 in AF647, clone B8.2C12, rat IgG1, BioLegend. Anti-CD103 in Biotin, clone 2E7, Armenian hamster IgG1, BioLegend. Anti-Streptavidin in BUV 395, BD. Anti-Ki67 in BV 480, clone B56, mouse IgG1, BD. Anti-TNF*α* in BV711, clone MP6-XT22, rat IgG1, BioLegend. Anti-INF*γ*BUV 737, clone XMG1.2, rat IgG1, BD. Anti-CD4 in BUV 496, clone GK1.5, rat IgG2b, BD. Anti-FOXP3 in PE, clone FJK-16s, rat IgG2a, ThermoFisher Scientific. Anti-EOMES in PE-eFluor610, clone Dan11mag, rat IgG2a, ThermoFisher Scientific. Anti-T-bet in PE-Cy7, clone eBio4B10, rat IgG1, ThermoFisher Scientific. Anti-CTLA4 in APC-AR700, clone UC10-4F10-11, armenian hamster IgG1, BD. Anti-CD24 in FITC, clone M1/69, rat IgG2b, BioLegend.

Flow cytometry data were compensated and exported using FlowJo software (version 10, TreeStar Inc.). The exported FCS files were normalized using Cyt3 MATLAB (version 2017b) and uploaded into Rstudio (R software environment, version 3.4.0). tSNE and FlowSOM algorithm mapping live cells from a pooled sample were performed as described^74^. CellCNN was run using default parameters, dividing data into training and validation steps as described^33^.

### Purification of CD39^+^PD-1^+^CD8^+^ T cells

Primary tumors were processed as described under “Flow Cytometry”. After the Percoll gradient, the leukocyte fraction was enriched for CD8^+^ T cells using anti-mouse CD8a microbeads (Miltenyi Biotec) and autoMACS equipment (Miltenyi Biotec) according to the manufacturer’s instructions. Subsequently, cells were stained for CD44, CD39, PD-1 and CD8 as described under “Flow Cytometry” and live, single CD44^+^CD39^+^PD-1^+^CD8^+^ T cells as well as other CD44^+^CD8^+^ T cells were sorted using an ARIA III Sorter (BD Biosciences).

### Adoptive transfer of CD39^+^PD-1^+^CD8^+^ T cells

Sorted cells were counted using trypan blue (Trypan Blue solution, Sigma Aldrich) to exclude dead cells. Twohundred-thousand CD44^+^CD39^+^PD-1^+^CD8^+^ T cells were injected immediately after 10^5^ 4T07-LZ cells. Injections were given intravenously.

### In vitro induction of senescence

Twenty-thousand 4T07 cells were plated in a 6-well plate. Subsequently, 75 ng/ml mouse IFNγ (R&D Systems) plus 5 ng/ml mouse TNF*α* (PeproTech) were added and cells were incubated at 37°C in a humidified atmosphere with 5% CO_2_ for 6 days as described. The proportion of senescent cells was determined by staining for *β*-galactosidase activity using Senescence *β*-Galactosidase Staining Kit (9860, Cell Signaling Technology) as described^42^.

### RNA-sequencing

#### Library preparation

The libraries were prepared following the Smart-seq2 protocol^75^. Five-hundred pg of cDNA from each sample were tagmented and amplified using Illumina Nextera XT kit. The resulting libraries were pooled, double-sided size selected (0.5x followed by 0.8x ratio using Beckman Ampure XP beads) and quantified using an Agilent 4200 TapeStation System. The pool of libraries was sequenced in an Illumina NovaSeq6000 sequencer (single-end 100 bp) with a depth of around 20 Mio reads per sample as a service by the Functional Genomics Center Zurich (FGCZ).

#### Data evaluation

The raw data generated by Illumina NovaSeq6000 sequencer were analyzed using SUSHI, a framework for analysis of NGS data developed by the FGCZ (Hatakeyama et al., 2016, Qi et al., 2017). Briefly, after quality control, the reads were aligned to a reference genome (Ensembl genome Mus_musculus. GRCm38 dated 2018.02.26) with STAR 2.7.3a. The software package EdgeR was used to detect differentially expressed genes. We applied a threshold of *p* < 0.01, FDR < 1% and a log fold-change of > 1.0 for upregulated genes and < -1.0 for downregulated genes. Unsupervised hierarchical clustering was done using the Ward2 method. Heatmaps were generated with the software FunRich V3.1.3.

Gene set enrichment analysis^43, 76^ was performed on a list of genes ranked from high to low estimated fold-change using the GSEA 4.0.3 Software (Broad Institute) with enrichment statistic classic and 1000 permutations.

### Patient material

Tumor tissues from 54 patients with breast cancer (Table 1) were collected at the University Hospital Zurich, Switzerland. Donors provided written, informed consent to tissue collection, analysis and data publication according to the Declaration of Helsinki. Law abidance was reviewed and approved by the ethics commission of the Canton Zurich (BASEC-Nr. 2018-02282 and KEK-ZH-2013-0584). Samples were numerically coded to protect donors’ rights to confidentiality and privacy.

### Immunofluorescence of human samples

Formalin-fixed paraffin-embedded samples were cut into 2 µm-thick sections and dried overnight at 55°C. Antigen retrieval and deparaffinization were performed in 1x Trilogy Solution (Cell Marque, 920P-06) for 15 minutes at 115°C. Staining was performed employing tyramide signal amplification (TSA) and the Opal^TM^ 7-color Manual IHC Kit (PerkinElmer). To prevent unspecific binding of antibodies, slides were incubated with 4% donkey serum in PBS (ANAWA BioWorld) in PBS/0.1% Triton X-100 for 15 minutes at 37°C. TSA staining protocol was performed as described^77^. Briefly, slides were incubated overnight at 4°C with primary antibody diluted in PBS/1% donkey serum/0.1% Triton X-100. Subsequently, slides were washed in PBS Triton X-100 0.1% and incubated with corresponding HRP-conjugated secondary antibodies. Following primary antibodies were used: Anti-CD39 (mouse monoclonal antibody, clone A1, BioLegend, 1:500), anti-PD-1 (rabbit monoclonal antibody, clone D4W2J BioConcept, 1:4000), anti-CD8 (mouse monoclonal antibody, clone RPA-T8, Cell Signaling, 1:2000), anti-PAN-Cytokeratin (rabbit polyclonal antibody, cat. no. H-240, Santa Cruz, 1:2000) and anti-EpCAM (rabbit monoclonal antibody, clone EPR20532-225, Abcam, 1:200). Following HRP-conjugated secondary antibodies were used: anti-mouse IgG (H+L) (donkey polyclonal antibody, 715-035-151 Jackson ImmunoResearch 1:500) and anti-rabbit IgG (H+L) (donkey polyclonal antibody, 715-035-152 Jackson ImmunoResearch 1:1000). Fluorescent signal was developed using the Opal^TM^ 7-color Manual IHC Kit. Finally, slides were washed, incubated with 0.5 µg/ml 4′,6-diamidine-2-phenylindole (DAPI; Invitrogen) for 5 min, washed again and mounted with ProlongDiamond medium (Invitrogen). The slides were scanned using the automated multispectral microscopy system Vectra 3.0 (PerkinElmer). An unstained slide was used to generate the spectral profile of autofluorescence in studied tissues. The Inform software (PerkinElmer) was used for (i) unmixing the spectra of individual fluorophores and autofluorescence, (ii) performing automated tissue and cell segmentation into tumor and non-tumor cells, and (iii) exporting single-cell intensity values for all fluorescent signals. Single-cell parameter files were transformed into .fcs files using R script (version 3.6.1) involving FlowCore and Biobase packages and were analyzed using FlowJo software to gate on CD39^+^PD-1^+^CD8^+^ T cell populations in tumor tissue category (Supplemental Figure 7F). Gates were set based on the intensity values identified in the fluorescent images. Cell density values were measured as number of cells/mm^2^ obtained as cell counts from the gated populations divided by the tumor tissue area measured in the corresponding patient. Samples were analyzed blinded.

### Statistical analysis

Data were analyzed using GraphPad Prism 8.0 for Windows or Mac, and RStudio (version 1.2.5019). For comparison of 2 experimental groups, 2-tailed Student’s t test with Welch’s correction was performed. More than 2 groups were compared using an ANOVA test with Bonferroni’s correction. Data following a logarithmic distribution (i.e. IVIS signal) were log transformed prior to analysis. The number of mice per experimental group was determined based on our previous experience with similar models. Every point represents one mouse. Unless stated otherwise, data are shown as mean ± SD. *p* < 0.05 is considered significant throughout. \**p* < 0.05, \*\**p* < 0.01, \*\*\**p* < 0.001, \*\*\*\**p* < 0.0001. CellCnn data were analyzed using the Kolmogorov–Smirnov two-sample statistical test. In addition, the population identified by the algorithm was compared in the two groups using the 2-tailed Student’s t test with Welch’s correction.

Density of CD39^+^PD-1^+^CD8^+^ T cells was correlated with the clinicopathological information of the patients using corresponding R packages. Disease-free survival was analyzed by Kaplan-Meier curve and log-rank test (Survminer and Survival R packages), as well as univariate and multivariate Cox regression (pROC, ROCR and Survival R packages). The threshold of CD39^+^PD-1^+^CD8^+^ or CD8^+^ T cell density was identified by the Receiver Operating Characteristics (ROC) curve analysis using 3-year disease-free survival for defining long- and short-term survivors. Following values apply: Intra-tumoral CD39^+^PD-1^+^CD8^+^ T cells, 4.20; extra-tumoral CD39^+^PD-1^+^CD8^+^ T cells, 8.43; total CD39^+^PD-1^+^CD8^+^ T cells, 5.71; intra-tumoral CD8^+^ T cells, 249.20; extra-tumoral CD8^+^ T cells, 1218.60; total CD8^+^ T cells, 562.29.

## Data deposition

https://www.ncbi.nlm.nih.gov/sra/PRJNA609233

## Materials Availability

All unique reagents generated in this study are available from the Lead Contact upon request.

## Legends to the supplemental figures

**Supplemental Figure S1. Circulating and disseminated cancer cells in mice with 4T1 or 4T07 orthotopic tumors.**

**(A)** Experimental design. 4T1 or 4T07 cells (10^5^) were injected into the mammary fat pad of female BALB/c mice. Analysis was performed 30 d later. **(B)** Growth curve of primary breast cancer. **(C)** Representative images of lungs at the endpoint stained with India ink to macroscopically visualize metastatic nodules. **(D)** The number of metastatic nodules in the lungs. **(E)** BALB/c mice were injected with 10^5^ 4T1 or 4T07 cells in the mammary fat pad and disseminated cancer cells (DCCs) were detected by colony-forming assay in the lungs 30 d later. DCCs were detected in all mice (n=5 per group). Representative images of colonies from cultured lung-digests are shown. **(F)** 4T07-mCh cells (10^5^) were injected into the mammary fat pad of female BALB/c mice. Analysis was performed 35 d later. Disseminated 4T07 cells were found in 5/5 mice and are shown in red, Ki67 in green, DAPI in blue. Representative pictures from a cohort of 5 mice. Scale bar: 20 µm.

Results are representative of 3 independent experiments.

**Supplemental Figure S2. 4T07 and 4T1 induce similar systemic inflammation.**

Female BALB/c mice were injected with 10^5^ 4T1 or 4T07 cells in the mammary fat pad. Control mice were injected with PBS. Analysis was performed at the indicated days after cancer cell injection. **(A)** Spleen weight. **(B)** Percentage circulating neutrophils of CD45^+^ cells. **(C-F)** Total number of cells per mg lung tissue: inflammatory monocytes (C), neutrophils (D), eosinophils (E), alveolar macrophages (F). **(G)** Female BALB/c mice were injected with 10^5^ 4T1 cells into the mammary fat pad. The plots show a representative gating strategy of lungs processed 20 d after tumor cell injection.

Each symbol represents an individual mouse. Five mice per group. ns = not significant (2-tailed Student’s t-test with Welch’s correction). The bar represents the mean ± SD.

**Supplemental Figure S3. 4T07 breast cancer elicits systemic protective immunity.**

**(A)** 4T07 cells (10^5^) or PBS were injected into the mammary fat pad of female BALB/c mice on d 0. On d 11, 3×10^5^ 4T07-LZ cells were injected i.v., and lung metastatic load (seeding) was quantified by bioluminescence 30 min and 3 h later. **(B)** Bioluminescence images. ns = not significant (2-tailed Student’s t-test with Welch’s correction). Each symbol represents an individual mouse. The bar represents the mean ± SD. Results are representative of 2 independent experiments. **(C)** Experimental design. 4T07 cells (10^5^) or PBS were injected into the mammary fat pad of female BALB/c mice on d 0. On d 11, 10^5^ 4T07-mCh cells were injected i.v. and analysis of lung metastatic load was performed on d 25. **(D)** Representative section of the lung at the endpoint. Upper panels: Disseminated 4T07 cells were detected as single, Ki67-negative cells in the lungs of 5 out of 5 mice. Disseminated 4T07 cells are shown in red, DAPI in blue. Lower panels: 4T07-metastatic nodules. Macro-metastasis was detected in 5 out of 5 mice. Scale bar indicates 20 µm. **(E)** Experimental design. 4T07 cells (10^5^) or PBS were injected into the mammary fat pad of female BALB/c mice on d 0. The tumor was resected on d 8; PBS-injected mice and one group of 4T07-injected mice underwent mock resection. On d 15, 10^5^ 4T07-mCh cells were injected i.v., and analysis of lung metastatic load was performed on d 30.

**(F)** Macroscopic pictures of lungs. Upper row: Metastatic nodules seen in magnifications of the circled areas. Middle and lower row: Absence of macroscopically visible metastatic nodules. **(G)** Representative example of disseminated 4T07 cells present in the lungs as single, Ki67-negative cells. Disseminated 4T07 cells are shown in red, Ki67 in green, DAPI in blue. Scale bar indicates 20 µm (upper panels) or 100 µm (lower panels). **(H)** Experimental design. 4T07 cells (10^5^) or PBS were injected into the mammary fat pad of female BALB/c mice on d 0. The tumor was resected on d 8 (PBS-injected mice underwent mock surgery) and 3×10^5^ 4T07-LZ cells were i.v. injected on d 15. The lung metastatic load was quantified on d 30. **(I)** Quantification of lung metastatic load by bioluminescence. \*\*\*\**p* < 0.0001, (2-tailed Student’s t-test with Welch’s correction). Each symbol represents an individual mouse. The bar represents the mean ± SD. Results are representative of 2 independent experiments. **(J)** Experimental design. 4T1 cells (10^5^), 4T07 cells (10^5^) or PBS were injected into the mammary fat pad of female BALB/c mice on d 0. On d 11, 3×10^5^ 4T1-LZ cells were injected and analysis of lung metastatic load was performed on d 25. **(K)** Quantification of lung metastatic load by bioluminescence. \**p* < 0.05, \*\*\*\**p* < 0.0001, ns = not significant (ANOVA with Bonferroni correction). Each symbol represents an individual mouse. The bar represents the mean ± SD.

**Supplemental Figure S4. Expression of selected markers by CD39^+^PD-1^+^CD8^+^ T cells.**

Female BALB/c mice were injected with 10^5^ 4T1 or 4T07 cells in the mammary fat pad and primary tumors were analyzed by flow cytometry 20 d later (*n*=4). **(A)** Gating strategy for Figure 3. **(B)** Representative examples of expression of selected markers on live, single CD45^+^ CD44^+^TCR*β*^+^CD11b^-^CD39^+^PD-1^+^CD8^+^ T cells. **(C)** Representative examples of IFNγ and TNF*α* production by CD45^+^CD44^+^TCR*β*^+^CD11b^-^CD39^+^PD-1^+^CD8^+^ T cells sorted from 4T07 tumors after in vitro stimulation with 100 ng/ml phorbol 12-myristate 13-acetate (PMA) plus 1 µg/ml ionomycin for 4 h at 37°C in the presence of GolgiPlug/GolgiStop.

**Supplemental Figure S5. IFNγ and TNF*α* induce senescence of 4T07 cells.** Twenty-thousand 4T07 cells were plated in a 6-well plate. Subsequently, 75 ng/ml mouse IFNγ plus 5 ng/ml mouse TNF*α* were added and cells were incubated for 6 days. **(A)** Proportion of senescent cells in control and IFNγ plus TNF*α*-exposed 4T07 cells. Each symbol represents an individual culture. \*\**p* < 0.01 (2-tailed Student’s t-test with Welch’s correction). The bar represents the mean ± SD. Results are representative of 2 independent experiments. **(B)** Representative examples of staining for SA-*β*-galactosidase activity (blue) as proxy for senescence. Left panel, medium. Right panel, IFNγ plus TNF*α*.

**Supplemental Figure S6. CD39^+^PD-1^+^CD8^+^ T cells in dormant and metastatic tumors have a similar transcription signature.**

**(A)** Hierarchical clustering of all genes by sorted CD44^+^CD39^+^PD-1^+^CD8^+^ and CD44^+^CD8^+^ T cells. **(B)** Volcano plot comparing transcripts in CD44^+^CD39^+^PD-1^+^CD8^+^ T cells sorted from 4T07 breast tumors with those sorted from 4T1 breast tumors. The red symbols represent transcripts that are significantly overexpressed in CD44^+^CD39^+^PD-1^+^CD8^+^ T cells from 4T07 breast tumors, whereas the blue symbols represent significantly under-expressed transcripts. **(C, D)** Venn diagram showing **(C)** shared or **(D)** significantly differentially expressed genes in CD44^+^CD39^+^PD-1^+^CD8^+^ and CD44^+^CD8^+^ T cells sorted from 4T1 and 4T07 breast cancers.

**Supplemental Figure S7. High density of intra-tumoral CD39^+^PD-1^+^CD8^+^ but not total CD8^+^ T cells correlate with disease-free survival in human luminal A and B breast cancer.**

**(A)** Disease-free survival of patients with luminal A, luminal B or triple-negative (TN) breast cancer. **(B)** Disease-free survival of 54 patients with high or low number of extra-tumoral CD39^+^PD-1^+^CD8^+^ T cells. **(C)** Disease-free survival of 54 patients with high or low number of intra-tumoral + extra-tumoral CD39^+^PD-1^+^CD8^+^ T cells. **(D)** Disease-free survival of 20 luminal A and 25 luminal B breast cancer patients with high or low number of intra-tumoral CD39^+^PD- 1^+^CD8^+^ T cells**. (E)** Disease-free survival of 20 luminal A and 25 luminal B breast cancer patients with high or low number of intra-tumoral CD8^+^ T cells. ns = not significant. **(F)** Representative example of a sample with high (upper panels) or low (lower panels) number of intra-tumoral CD39^+^PD-1^+^CD8^+^ T cells. Single-cell parameter files were transformed into .fcs files using R studio and were analyzed using FlowJo software. Values of number of cells/mm^2^ were obtained by using the values of total counts obtained from the FlowJo analysis and the measurements of tumor tissue area obtained from the inform analysis. The threshold for separating patients with high and low CD39^+^PD-1^+^CD8^+^ T cell densities was defined using ROC curve analysis.

