## Supplemental Figures for "CD39^+^PD-1^+^CD8^+^ T cells mediate metastatic dormancy in breast cancer"

### Supplemental Figure 1

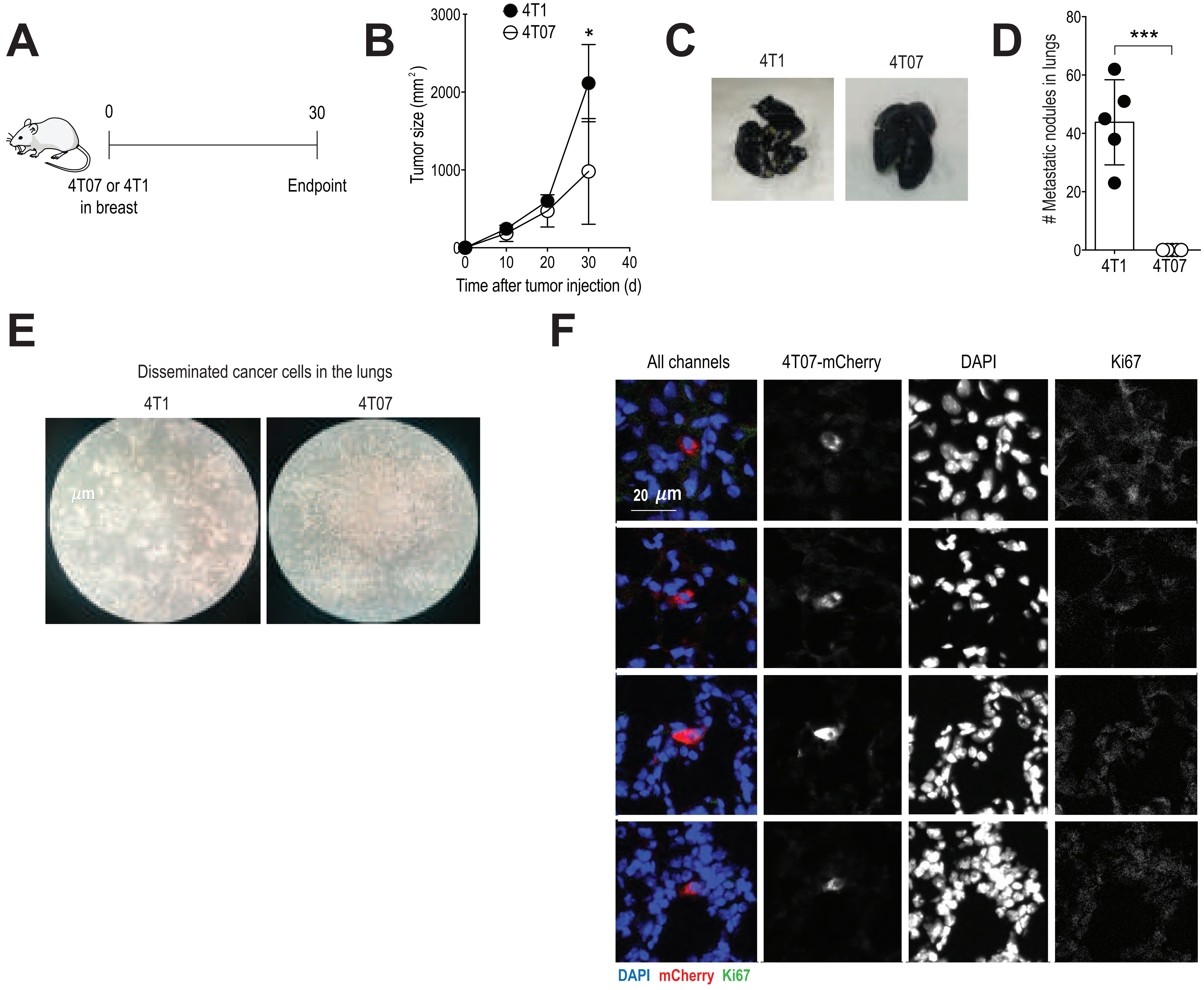

### Supplemental Figure 2

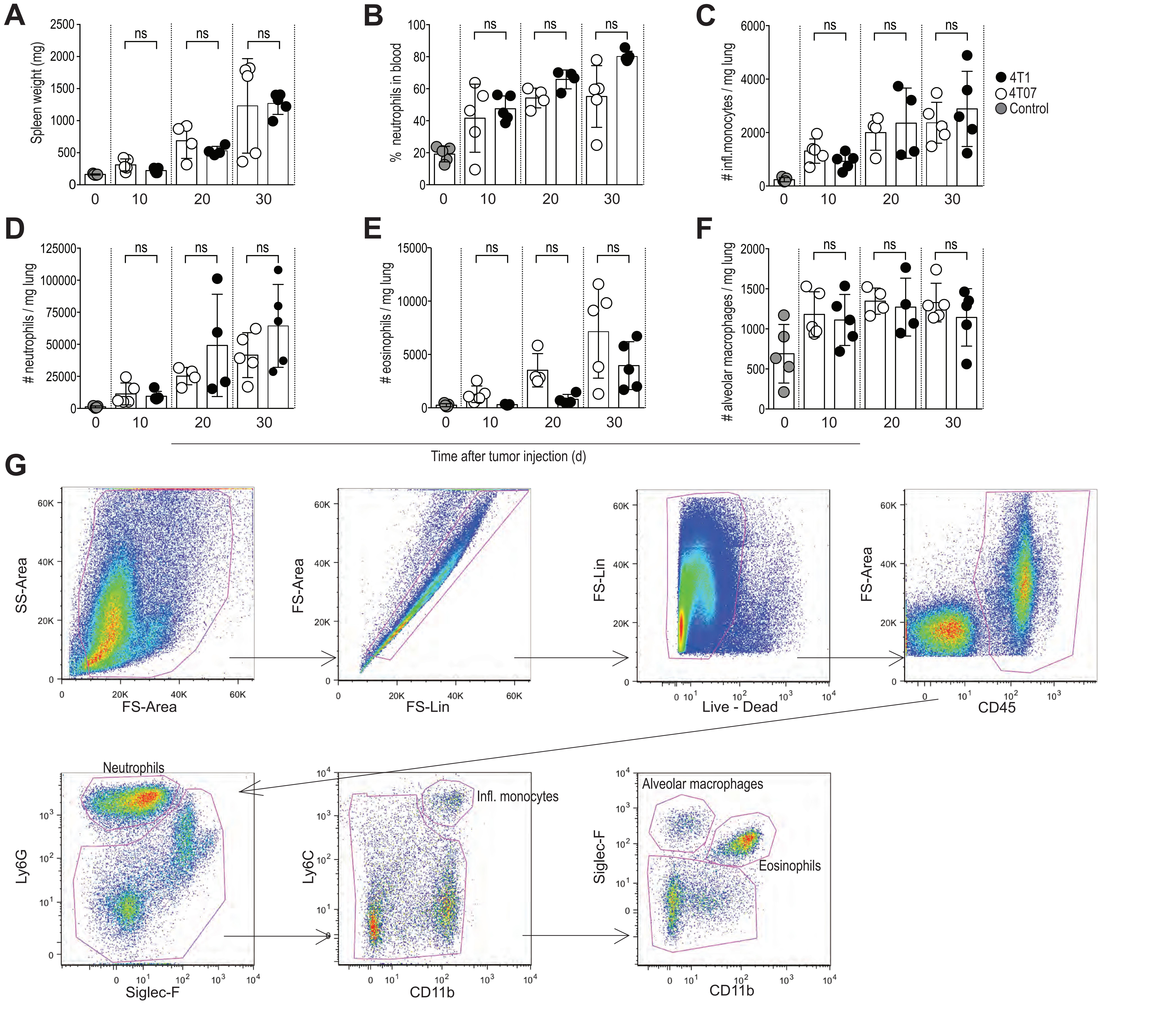

### Supplemental Figure 3

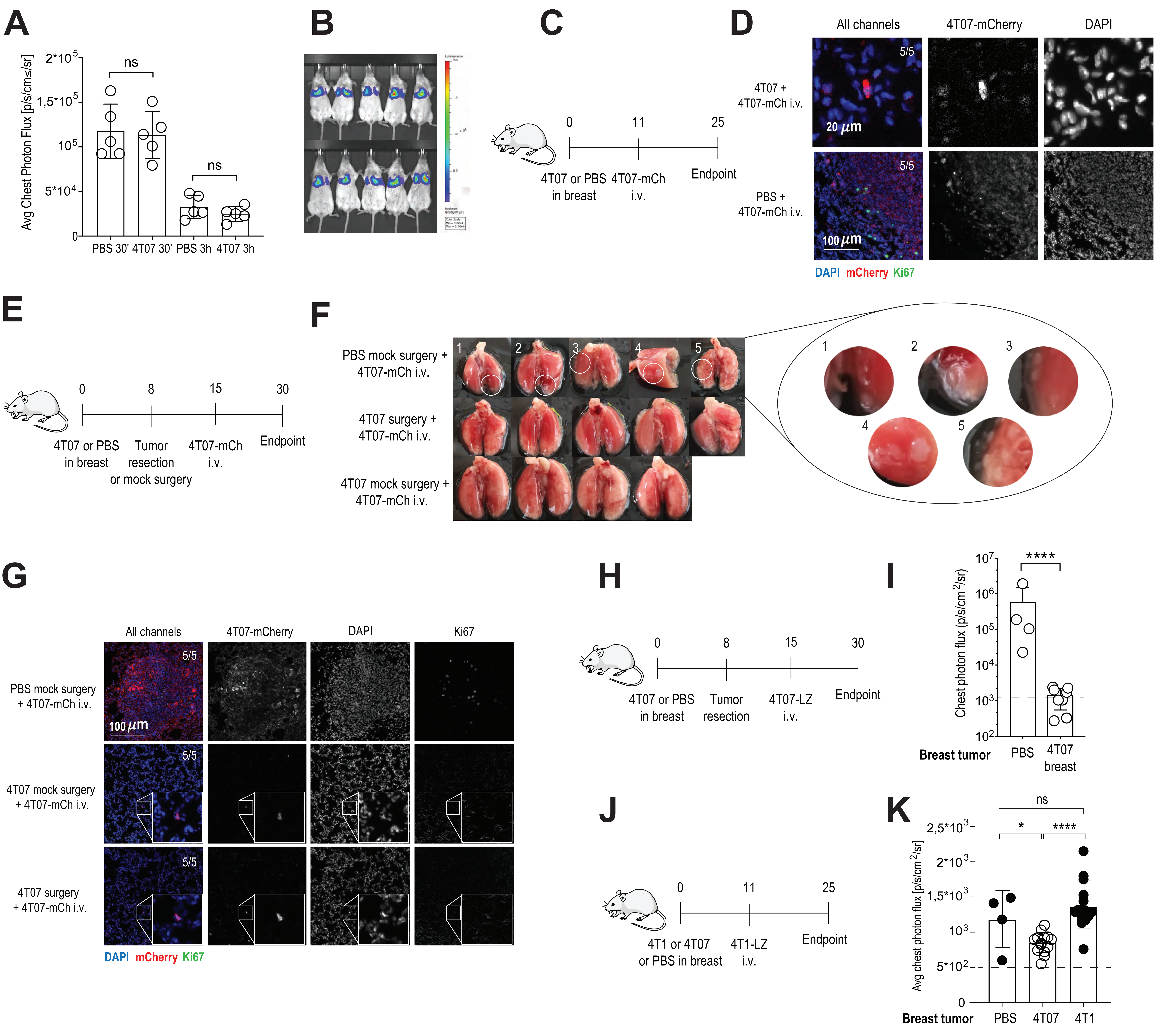

### Supplemental Figure 4

**A**

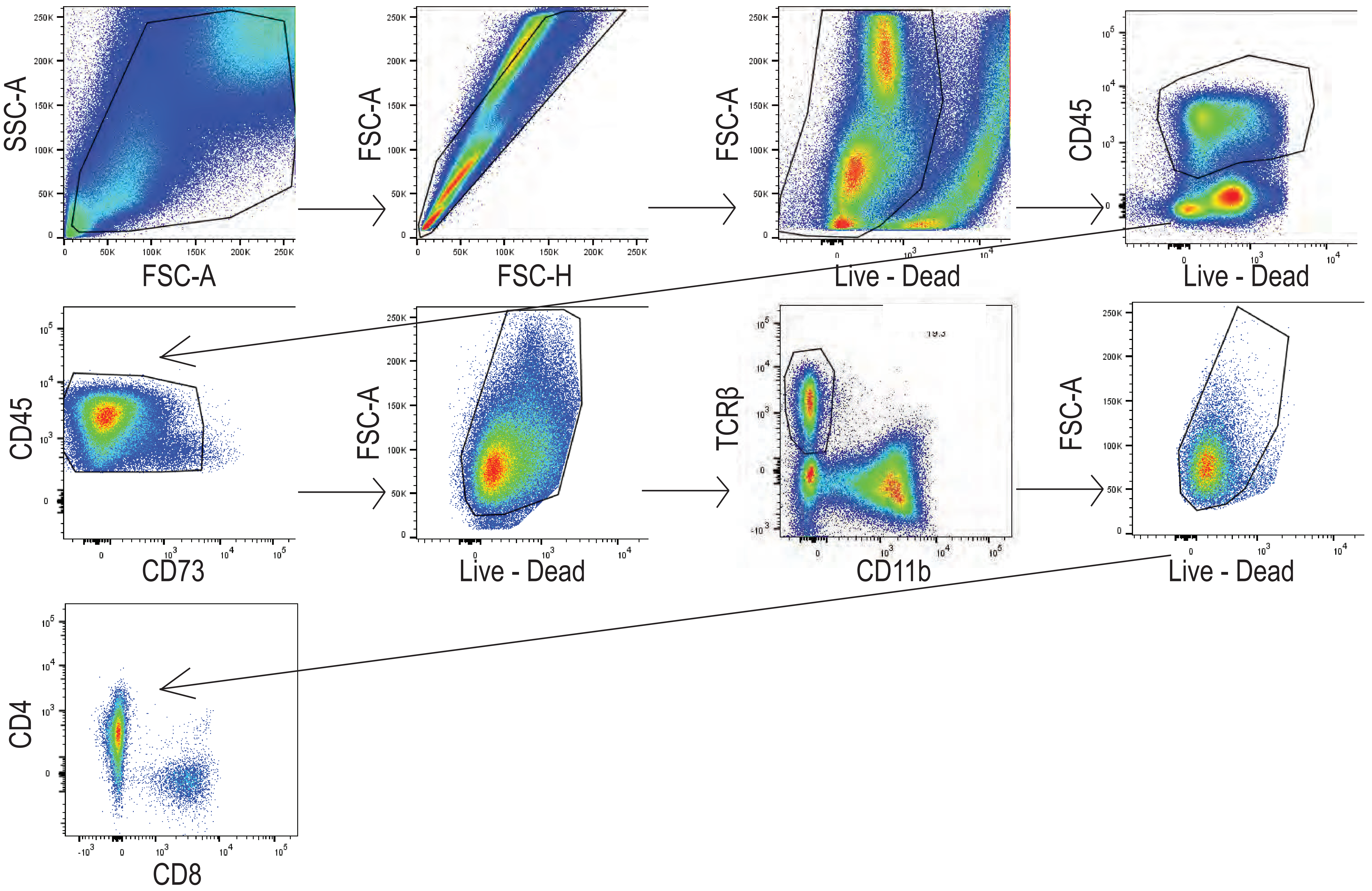


**B**

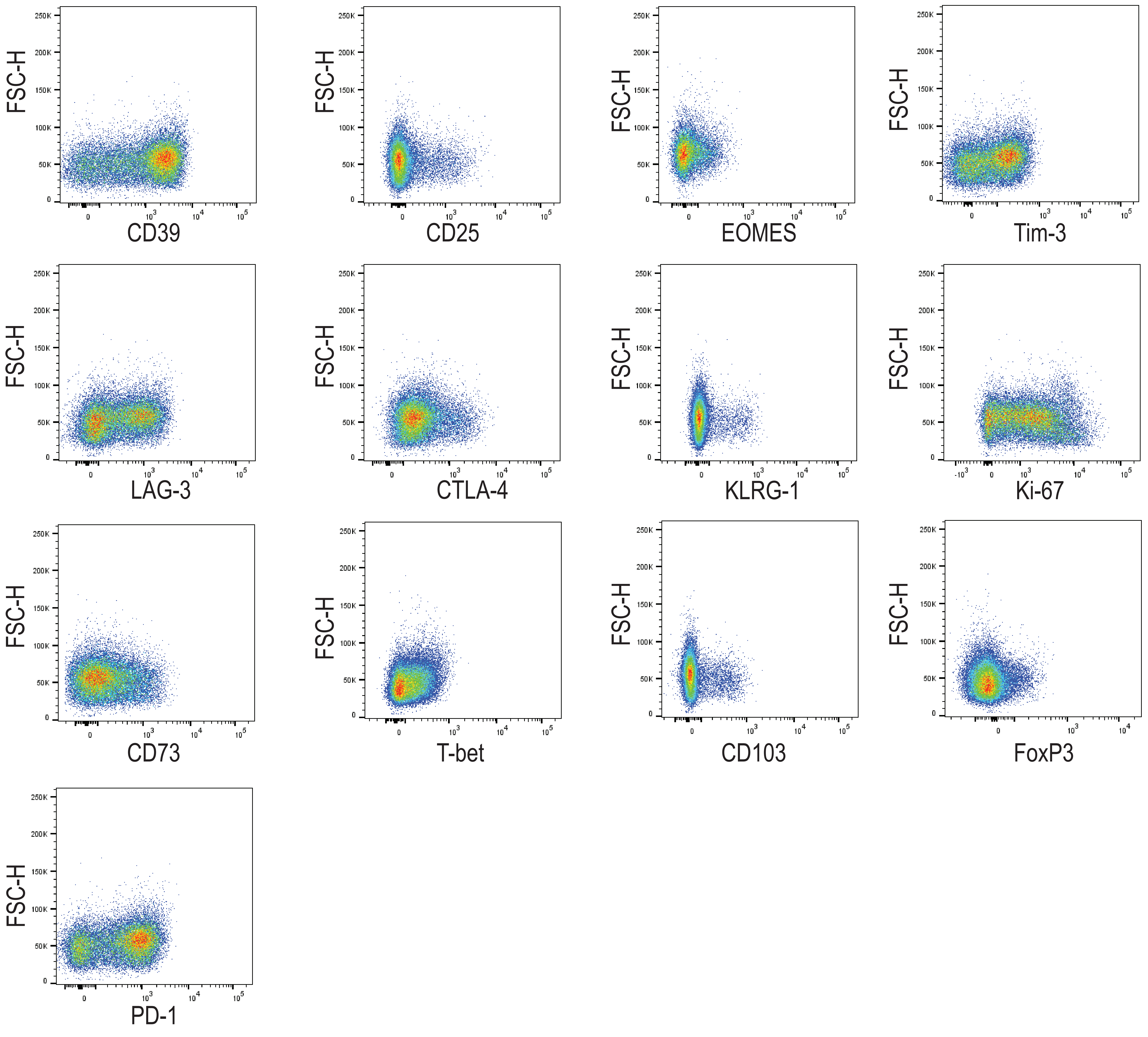


**C**

CD39⁺PD-1⁺CD8⁺ T cells

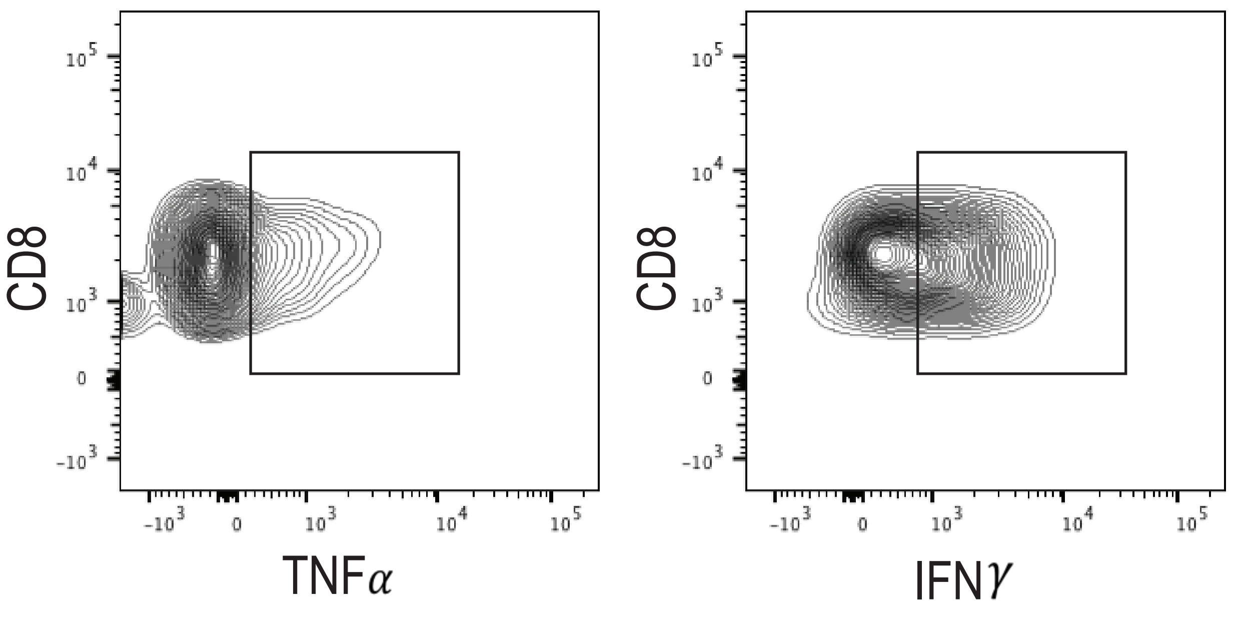

### Supplemental Figure 5

A

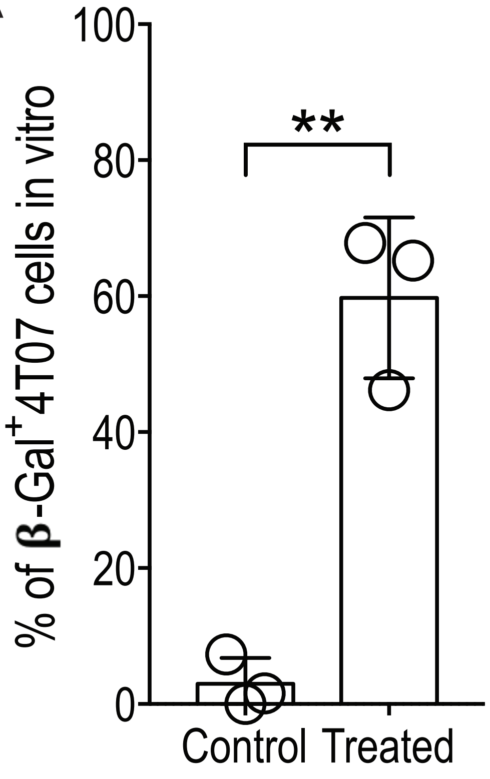


B

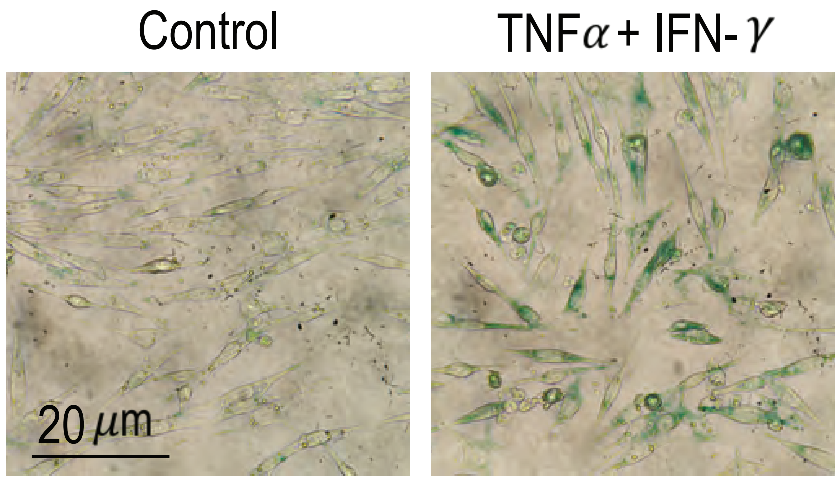

### Supplemental Figure 6

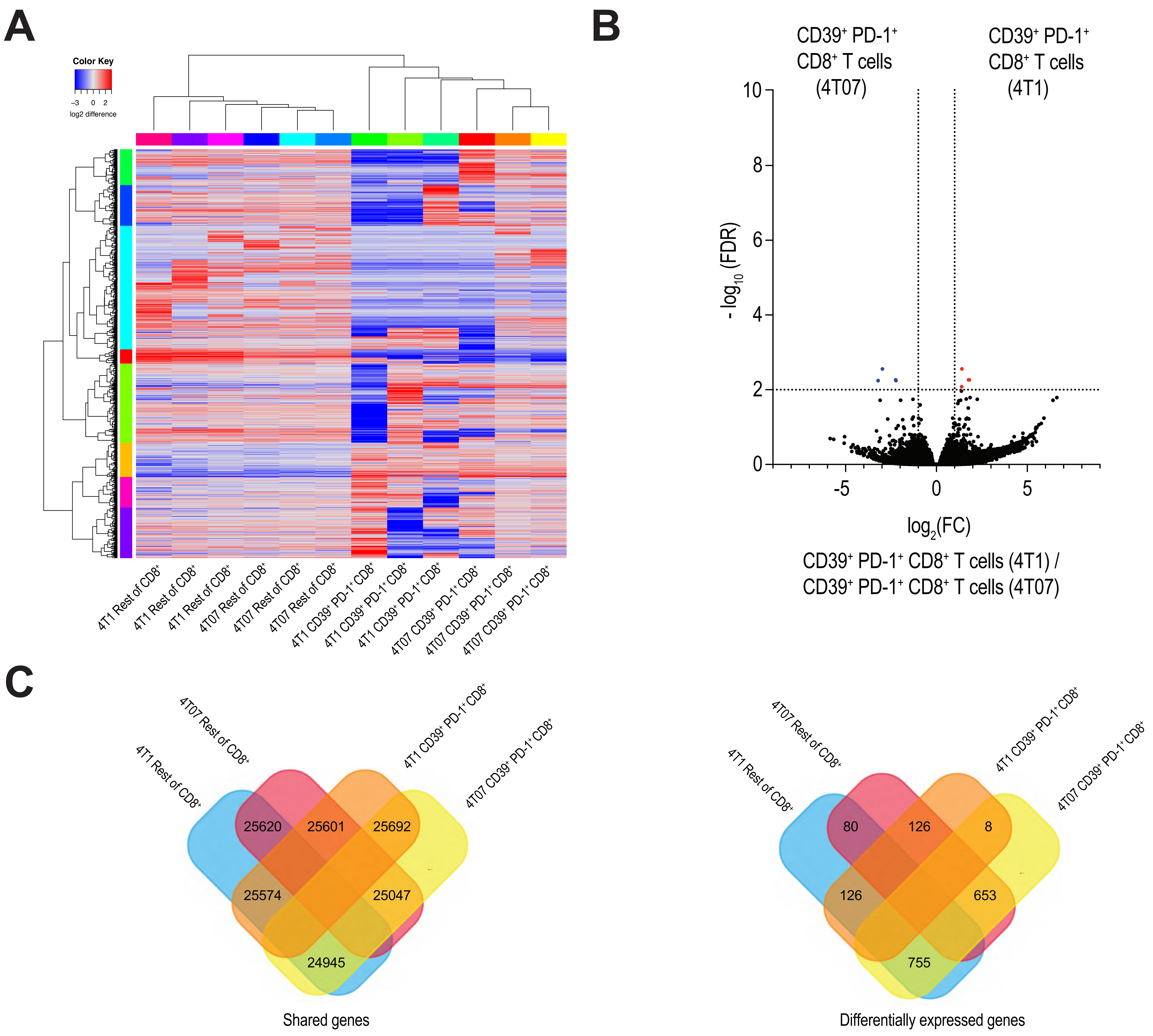

### Supplemental Figure 7

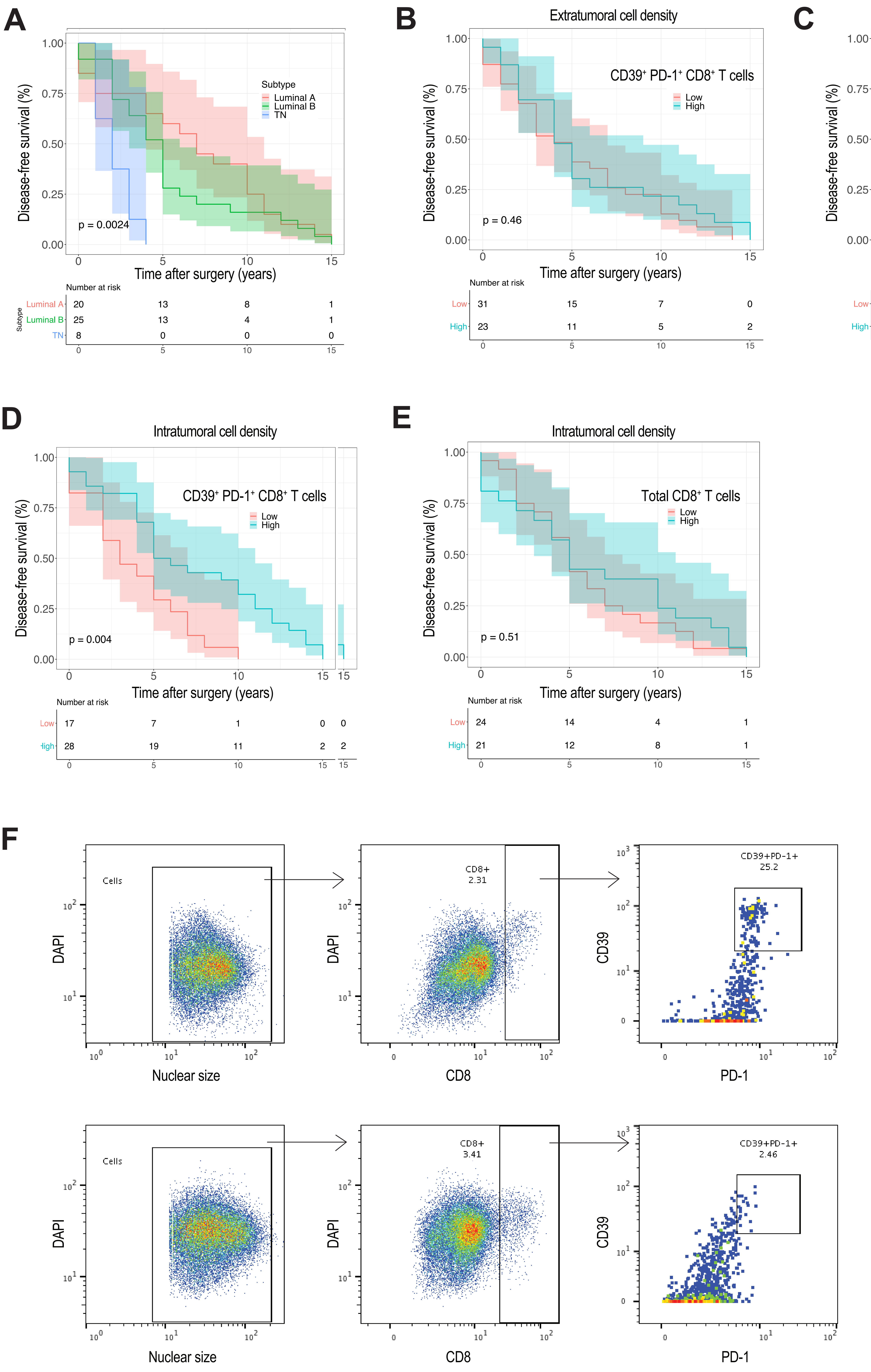

| Clinicopathological parameters n=54 | | | |
|---|---|---|---|
| | N° | % | Univariate-Cox *p* value |
| **Age at initial diagnosis** | | | 0.14 |
| Age < 55 | 27 | 50 | |
| Age ≥ 55 | 27 | 50 | |
| **Tumor (T) stage** | | | 0.0049 |
| T1 | 18 | 33.5 | |
| T2 | 29 | 53.5 | |
| T3 | 4 | 7 | |
| T4 | 3 | 6 | |
| **Nodal (N) stage** | | | 0.64 |
| Negative | 23 | 42.5 | |
| Positive | 31 | 57.5 | |
| **Tumor grade** | | | 0.074 |
| Grade 1 | 3 | 5.5 | |
| Grade 2 | 21 | 39 | |
| Grade 3 | 30 | 55.5 | |
| **ER status** | | | 0.00077 |
| Negative | 9 | 16.5 | |
| Positive | 45 | 83.5 | |
| **PR status** | | | 0.53 |
| Negative | 15 | 28 | |
| Positive | 39 | 72 | |
| **Her2 status** | | | 0.82 |
| Negative | 37 | 66.5 | |
| Positive | 17 | 31.5 | |
| **Molecular subtype** | | | 0.0013 |
| Luminal A | 20 | 37 | |
| Luminal B | 25 | 46 | |
| Her2 | 1 | 2 | |
| TN | 8 | 15 | |
| **Density of intratumoral $CD39^{+}PD\text{-}1^{+}CD8^{+}$ T cells** | NA | NA | Multivariate-Cox: Not independent |
